# Transcriptional Cartography Integrates Multiscale Biology of the Human Cortex

**DOI:** 10.1101/2022.06.13.495984

**Authors:** Konrad Wagstyl, Sophie Adler, Jakob Seidlitz, Simon Vandekar, Travis T. Mallard, Richard Dear, Alex R. DeCasien, Theodore D. Satterthwaite, Siyuan Liu, Petra E. Vértes, Russell T. Shinohara, Aaron Alexander-Bloch, Daniel H. Geschwind, Armin Raznahan

**Affiliations:** Wellcome Centre for Human Neuroimaging, University College London, London, UK; UCL Great Ormond Street Institute for Child Health, 30 Guilford St, Holborn, London WC1N 1EH; Department of Psychiatry, University of Pennsylvania, Philadelphia, PA 19104; Department of Child and Adolescent Psychiatry and Behavioral Science, The Children’s Hospital of Philadelphia, Philadelphia, PA 19104; Department of Biostatistics, Vanderbilt University, Nashville, Tennessee, USA; Psychiatric and Neurodevelopmental Genetics Unit, Center for Genomic Medicine, Massachusetts General Hospital, Boston, MA, USA; Department of Psychiatry, Harvard Medical School, Boston, MA, USA; Department of Psychiatry, University of Cambridge, Cambridge, CB2 0SZ, UK; Section on Developmental Neurogenomics, Human Genetics Branch, National Institute of Mental Health, Bethesda, MD, USA; Lifespan Informatics and Neuroimaging Center, University of Pennsylvania School of Medicine, Philadelphia, PA, 19104; Penn Statistics in Imaging and Visualization Center, Department of Biostatistics, Epidemiology, and Informatics, Perelman School of Medicine, University of Pennsylvania, Philadelphia, PA, USA; Center for Autism Research and Treatment, Semel Institute, Program in Neurogenetics, Department of Neurology, and Department of Human Genetics, David Geffen School of Medicine, University of California Los Angeles, Los Angeles, CA, US

## Abstract

The cerebral cortex underlies many of our unique strengths and vulnerabilities - but efforts to understand human cortical organization are challenged by reliance on incompatible measurement methods at different spatial scales. Macroscale features such as cortical folding and functional activation are accessed through spatially dense neuroimaging maps, whereas microscale cellular and molecular features are typically measured with sparse postmortem sampling. Here, we integrate these distinct windows on brain organization by building upon existing postmortem data to impute, validate and analyze a library of spatially dense neuroimaging-like maps of human cortical gene expression. These maps allow spatially unbiased discovery of cortical zones with extreme transcriptional profiles or unusually rapid transcriptional change which index distinct microstructure and predict neuroimaging measures of cortical folding and functional activation. Modules of spatially coexpressed genes define a family of canonical expression maps that integrate diverse spatial scales and temporal epochs of human brain organization - ranging from protein-protein interactions to large-scale systems for cognitive processing. These module maps also parse neuropsychiatric risk genes into subsets which tag distinct cyto-laminar features and differentially predict the location of altered cortical anatomy and gene expression in patients. Taken together, the methods, resources and findings described here advance our understanding of human cortical organization and offer flexible bridges to connect scientific fields operating at different spatial scales of human brain research.

## Introduction

The human cerebral cortex is an astoundingly complex structure that underpins many of our distinctive facilities and vulnerabilities(Geschwind and Rakic, 2013). Achieving a mechanistic understanding of cortical organization in health and disease requires integrating information across its many spatial scales: from macroscale cortical folds and functional networks(Glasser et al., 2016) to the gene expression programs that reflect microscale cellular and laminar features(Hawrylycz et al., 2012; Kelley et al., 2018). However, a hard obstacle to this goal is that our measures of the human cortex at macro- and microscales are fundamentally mismatched in their spatial sampling. Macroscale measures from in vivo neuroimaging provide spatially dense estimates of structure and function, but microscale measures of gene expression are gathered from spatial discontinuous postmortem samples that have so far only been linked to macroscale features using methodologically-imposed cortical parcellations(Hansen et al., 2021; Larivière et al., 2021; Seidlitz et al., 2020). Consequently, local transitions in human cortical gene expression remain uncharacterized and unintegrated with the spatially fine-grained topographies of human cortical structure and function that are revealed by in vivo neuroimaging(Gryglewski et al., 2018; Markello et al., 2021). Finding a way to bridge this gap would not only enrich both our micro- and macro-scale models of human cortical organization, but also provide an essential framework for translation across traditionally siloed scales of neuroscientific research.

Here, we use spatially sparse postmortem data from the Allen Human Brain Atlas [AHBA(Hawrylycz et al., 2012)] to generate spatially dense cortical expression maps (DEMs) for 20,781 genes in the adult brain, with accompanying DEM reproducibility scores to facilitate wider usage. These maps allow a fine-grained transcriptional cartography of the human cortex, which we integrate with diverse genomic, histological and neuroimaging resources to shed new light on several fundamental aspects of human cortical organization in health and disease. First, we show that DEMs can recover canonical gene expression boundaries from in situ hybridization (ISH) data, predict previously unknown expression boundaries and align with regional differences in cortical organization from several independent data modalities. Second, by focusing on the local transitions in gene expression which are captured by DEMs, we reveal a close spatial coordination between molecular and functional specializations of the cortex, and establish that the spatial orientation of cortical folding and function at macroscale is aligned with local tangential transitions in cortical gene expression. Third, by defining and annotating gene co-expression modules across the cortex at multiple scales we systematically link macroscale measures of cortical structure and function in vivo, to postmortem markers of cortical lamination, cellular composition and development from early fetal to late adult life. Finally, as a proof-of-principle, we use this novel framework to secure a newly-integrated multiscale understanding of atypical brain development in autism spectrum disorder (ASD).

The tools and results from this analysis of the human cortex - which we collectively call Multiscale Atlas of Gene expression for Integrative Cortical Cartography (MAGICC) - open up an empirical bridge that can now be used to connect cortical models (and scientists) that have so far operated at segregated spatial scales. To this end, we share: (i) all gene-level DEMs and derived transcriptional landscapes in neuroimaging-compatible files for easy integration with in vivo macroscale measures of human cortical structure and function; and (ii) all gene sets defining spatial subcomponents of cortical transcription for easy integration with any desired genomic annotation (https://github.com/kwagstyl/magicc).

## Results

### Creating and benchmarking spatially dense maps of human cortical gene expression

To create a dense transcriptomic atlas of the cortex, we used AHBA microarray measures of gene expression for 20,781 genes in each of 1304 cortical samples from six donor left cortical hemispheres (**Methods, Table S1**). We extracted a model of each donor’s cortical sheet by processing their brain MRI scan, and identified the surface location (henceforth “vertex”) of each postmortem cortical sample in this sheet (**Methods**, **Fig 1a**). For each gene, we then propagated measured expression values into neighboring vertices using nearest-neighbor interpolation followed by smoothing (**Methods**, **Fig 1b,c**). Expression values were scaled across vertices and these vertex-level expression maps were averaged across donors to yield a single dense expression map (DEM) for each gene - which provided estimates of expression at ∼ 30,000 vertices across the cortical sheet (e.g. DEM for PVALB upper panel **Fig 1d**). These fine-grained vertex-level expression measures also enabled us to estimate the orientation and magnitude of expression change for each gene at every vertex (e.g. dense expression change map for PVALB, lower panel **Fig 1d**)

**Figure 1.**
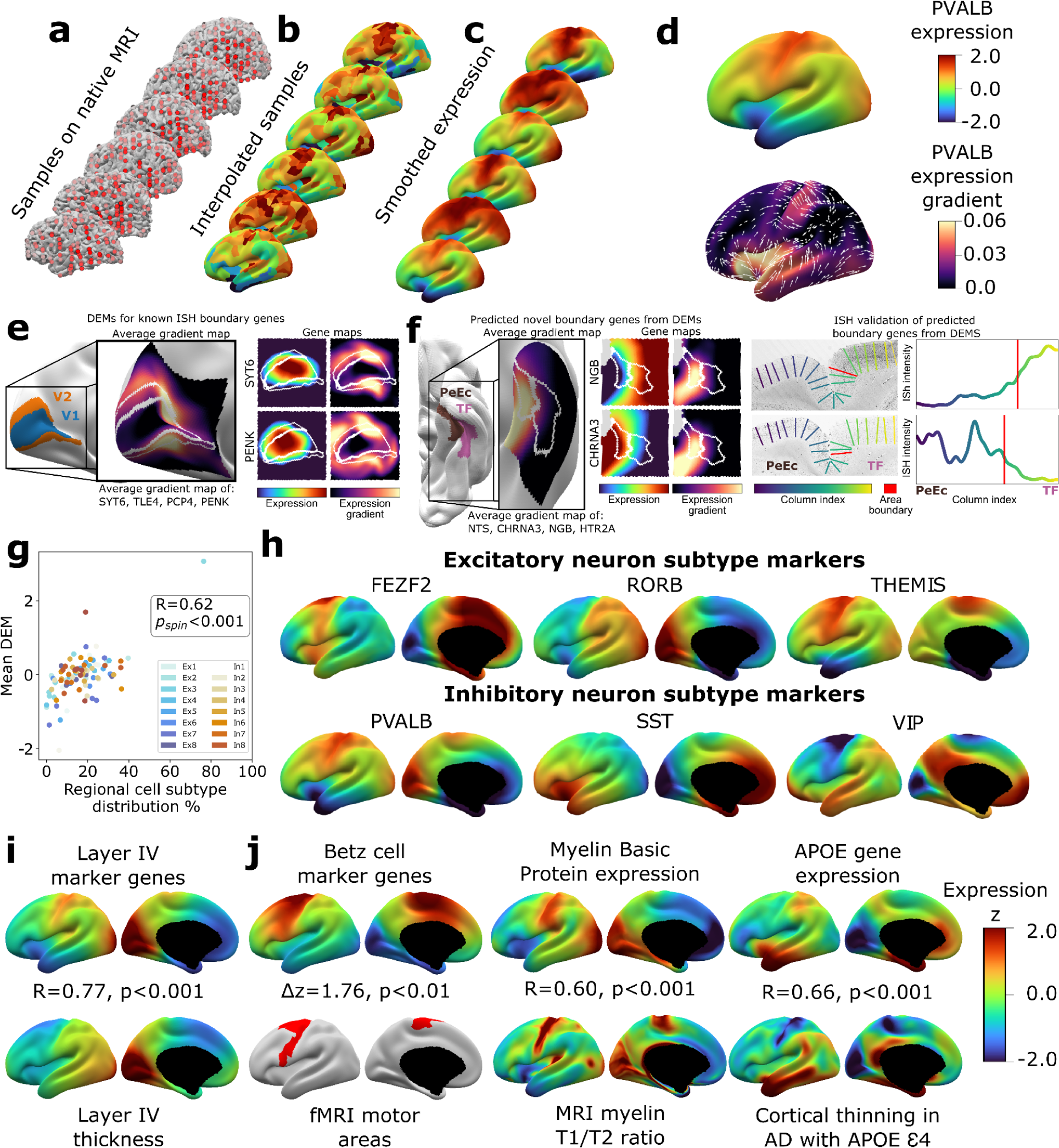
Creating and Benchmarking Spatial Dense Gene Expression Maps in the Human Cortex. **a**, Spatially discontinuous Allen Human Brain Atlas (AHBA) microarray samples (red points) were aligned with MRI-derived cortical surface mesh reconstructions. **b**, AHBA vertex expression values were propagated using nearest-neighbor interpolation and subsequently smoothed **(c)**. **d,** Subject-level maps were z-normalized and averaged to generate a single reference dense expression map (DEM) for each gene, as well as the associated expression gradient map (shown here for PVALB: top and bottom, respectively). **e,** DEMs can recover known expression boundaries in ISH data. Four canonical V1 area markers (Zeng et al., 2012 Cell) show a significantly sharp DEM expression gradient at the V1/V2 boundary (insert cortical map and **Fig S2a,b**), which is also evident in all four individual gene DEMs and DEM gradients (SYT6, PENK and **Fig S2c**). **f**, DEMs can discover previously unknown expression boundaries. Genes with high DEM gradients across the PeEc (parahippocampal) and TF (fusiform) gyri (inset cortical map) were validated in ISH data - showing sharp expression changes in both directions at this boundary (CHRNA3, NGB and **Fig Sd-f**).**g**, Illustrative comparisons of selected DEMs against regional variation in microscale measures of cellular composition: scatterplot showing the global correlation of regional cellular proportions from single nucleus RNAseq (snRNAseq) across 16 cells and 6 regions(Lake et al., 2016) with DEM values for corresponding cell-type marker genes (R=0.48, p_spin_<0.001, excluding Ex3-V1 and In8-BA10 outlier samples). **h**, DEMs for markers of 6 neuronal subtypes (3 excitatory: FEZF2, RORB, THEMIS, 3 inhibitory: PVALB, SST, VIP) based on recently validated subtype marker genes(Bakken et al., 2021; Hodge et al., 2019)**i,** Illustrative comparison of layer IV marker DEMs with corresponding mesoscale cortical measure of layer IV thickness from a 20μm 3D histological atlas of cortical layers. **j,** Illustrative comparisons of selected DEMs with corresponding macroscale cortical measures from independent neuroimaging markers.

We assessed the reproducibility of DEMs by repeating the above process (**Fig 1**) after repeatedly splitting the donors into non-overlapping groups of varying size, and using learning curve analyses to estimate the DEM reproducibility achieved by our full set of 6 donors. For cortically expressed genes (**Methods, Table S2**), the average reproducibility of gene expression maps was r_gene_=0.58 (correlation of expression values for a gene across vertices), and the average reproducibility of ranked gene expression at each vertex was r_vertex_=0.63 (correlation of expression values at a vertex across genes) (**Fig S1c-d**). These estimates were both substantially lower for genes not reported to be cortically expressed in the independent Human Protein Atlas (r_gene_=0.34, t=37.6, p<0.001 and r_vertex_=0.39, t=273.6, p<0.001, respectively, **Methods, Table S2**). Genes without recorded cortical expression were 3-fold enriched (p=0) amongst the 9,647 genes with estimated DEM reproducibility values of r <0.5). Regional differences in the density of postmortem sampling in the AHBA did not influence DEM reproducibility or the magnitude of local expression change captured by DEMs (**Methods, Fig S1h**). Thus, remedying the current lack of any spatially dense gene expression maps in the human cortex, we provide DEMs (and accompanying dense expression change maps) for 20,781 genes, and establish that >11k of these DEMs show a spatial reproducibility score of r_gene_>0.5 between sets of unrelated individuals. Gene-level DEM reproducibility scores allow future users to filter on this feature as desired, and we establish that key analytic outputs from DEMs (see below) show good reproducibility between unrelated individuals and can be recovered at different DEM reproducibility filters.

Given that DEMs were generated by interpolating expression values between sampled regions, we assessed if DEMs could recover sharp local microscale transitions in gene expression that could theoretically be obscured by interpolation. Of the very few such transitions that have been verified by ISH in humans, the best-established occurs between occipital areas V1 and V2(Zeng et al., 2012). All four genes known to show a sharp V1/V2 expression boundary across layers by ISH - SYT6, TLE4, PCP4, PENK - exhibited qualitatively and quantitatively sharp expression transitions at the V1/V2 boundary in their DEMs (**Fig 1e, Fig S2a-d**). Motivated by this validation, we next asked if DEMs could identify previously unknown expression boundary markers in the human cortex. To achieve this, we took advantage of extensive existing ISH data between parahippocampal (area PeEc) and fusiform gyri (area TF). We ranked genes by the magnitude of their expression gradient between these cortical regions in DEMs (**Methods**), and identified 4 genes with sharp expression transitions predicted by DEMs - NGB,HTR2A, (TF>PeEc) and NTS, CHRNA3 (PeEc>TF) - for which independent ISH data were available. Expression profiling in ISH slabs verified the existence of sharp expression transition for all four genes (**Fig 1f, Fig S2e-g**). As the V1/V2 and the PeEc/TF boundaries both involve transitions between classical laminar types in cortical regions with highly conserved anatomical patterning(von Economo and Koskinas, 1925), we also tested if DEMs could recover expression boundaries in more variable and uniformly laminated association cortex(Ronan and Fletcher, 2015). No such expression boundaries have been described in humans by ISH, but there are reports of sharp expression boundaries between frontal areas 44 and 45b for several genes in non-human primates: SCN1B, KCNS1, TRIM55(Chen et al., 2022). These genes also exhibited high DEM gradients at the boundary between human frontal areas 44 and 45 (**Fig S2h-j**). Taken together, these observations demonstrate the capacity of DEMs to resolve sharp expression transitions and indicate that DEMs can be used to help target prospective post mortem validation of new expression boundaries in humans.

To benchmark and illustrate the use of DEMs to capture cortical features across contrasting spatial scales, we drew on selected micro- and macro- and macroscale cortical measures that DEMs should align with based on known biological processes (**Fig 1g-j, Methods**). To assess if DEMs could recover microscale differences in cellular patterning across the cortical sheet, we considered the ground truth of neuronal cell-type proportions as measured by single nucleus RNAseq (snRNAseq) across 6 different cortical regions(Lake et al., 2016). We observed a strong spatial correlation (r=0.6, p_spin_<0.001) between regional marker gene expression in DEMs and regional proportions of their corresponding neuronal subtypes from snRNAseq (**Fig 1g, Methods**). **Fig 1h** shows example marker gene DEMs for 6 canonical neuronal subtypes: 3 excitatory (FEZF2, RORB, THEMIS) and 3 inhibitory (PVAL, SST, VIP)(Bakken et al., 2021; Hodge et al., 2019). Next, to assess if DEMs could recover regional variation in the mesoscale feature of cortical layering, we tested and verified that regional variation in the average DEM for layer IV marker genes(He et al., 2017; Maynard et al., 2021; Zeng et al., 2012) was highly correlated with regional variation in layer IV thickness as determined from a 3D histological atlas of cortical layers(Wagstyl et al., 2020) (**Fig 1i**). Finally, we asked if DEMs could recover spatially-dense measures of regional variation across the cortical sheet as provided by neuroimaging data, and found that maps from diverse measurement modalities showed strong and statistically-significant spatial correlations with their corresponding DEM(s) relative to a null distribution based on random “spinning” of maps(Alexander-Bloch et al., 2018) (**Fig 1j, Methods,** all p_spin_<0.01): (i) areas of cortex activated during motor fMRI tasks in humans(Glasser et al., 2016) vs. the average DEM for canonical cell markers of large pyramidal neurons (Betz cells) found in layer V of the motor cortex that are the outflow for motor movements(Bakken et al., 2021), (ii) an *in vivo* neuroimaging marker of cortical myelination (T1/T2 ratio(Glasser and Van Essen, 2011)) vs. the Myelin Basic Protein DEM, which marks myelin, and (iii) the degree of in vivo regional cortical thinning by MRI in Alzheimer disease patients who have at least one APOE E4 variant(Gutiérrez-Galve et al., 2009; LaMontagne et al., 2019) vs. the APOE DEM (thinning map generated from 119 APOE E4 patients and 633 controls structural MRI (sMRI) scans as detailed in **Methods**), testing the hypothesis that higher regional APOE expression will result in greater cortical atrophy in individuals with the APOE E4 risk allele. Collectively, the above tests of reproducibility (**Fig S1**) and convergent validity (**Fig 1e-j**) supported use of DEMs for downstream analyses.

### Defining and surveying the human cortex as a continuous transcriptional terrain

As an initial summary view of transcriptional patterning in the human cortex, we first averaged all 20,781 DEMs to represent the cortex as a single continuous transcriptional terrain, where altitude encodes the transcriptional distinctiveness (TD) of each cortical point (vertex) relative to all others (TD = mean(abs(z_exp_)), **Figure 2a**, **Sup Movie 1**). This terrain view revealed 6 statistically-significant TD peaks (**Methods**, **Fig. 2a,b**) which recover all major archetypal classes of the mammalian cortex as defined by classical studies of laminar and myelo-architecture, connectivity, and functional specialization(Mesulam, 1998) encompassing: primary visual (V1), somatosensory [Brodmann area (BA(Brodmann, 1909)) 2], and motor cortex (BA 4), as well limbic [temporal pole centered on dorsal temporal area G (TGd(von Economo and Koskinas, 1925)), ventral frontal centered in orbitofrontal cortex (OFC)] and heteromodal association cortex (BA 9-46d). Of note, our agnostic parcellation of all TD peak vertices by their ranked gene lists (**Methods**) perfectly cleaved BA2 and BA4 along the central sulcus - despite there being no representation of this macroanatomical landmark in DEMs. The TD map observed from the full DEMs library was highly stable between all disjoint triplets of donors (**Methods**, **Fig S3a**, median cross-vertex correlation in TD scores between triplets r=0.77) and across library subsets at all deciles of DEM reproducibility (**Methods**, **Fig S3b**, cross-vertex correlation in TD scores r>0.8 for the 3rd-10th deciles), but was not recapitulated in spun null datasets (**Fig S3c**).

**Figure 2.**
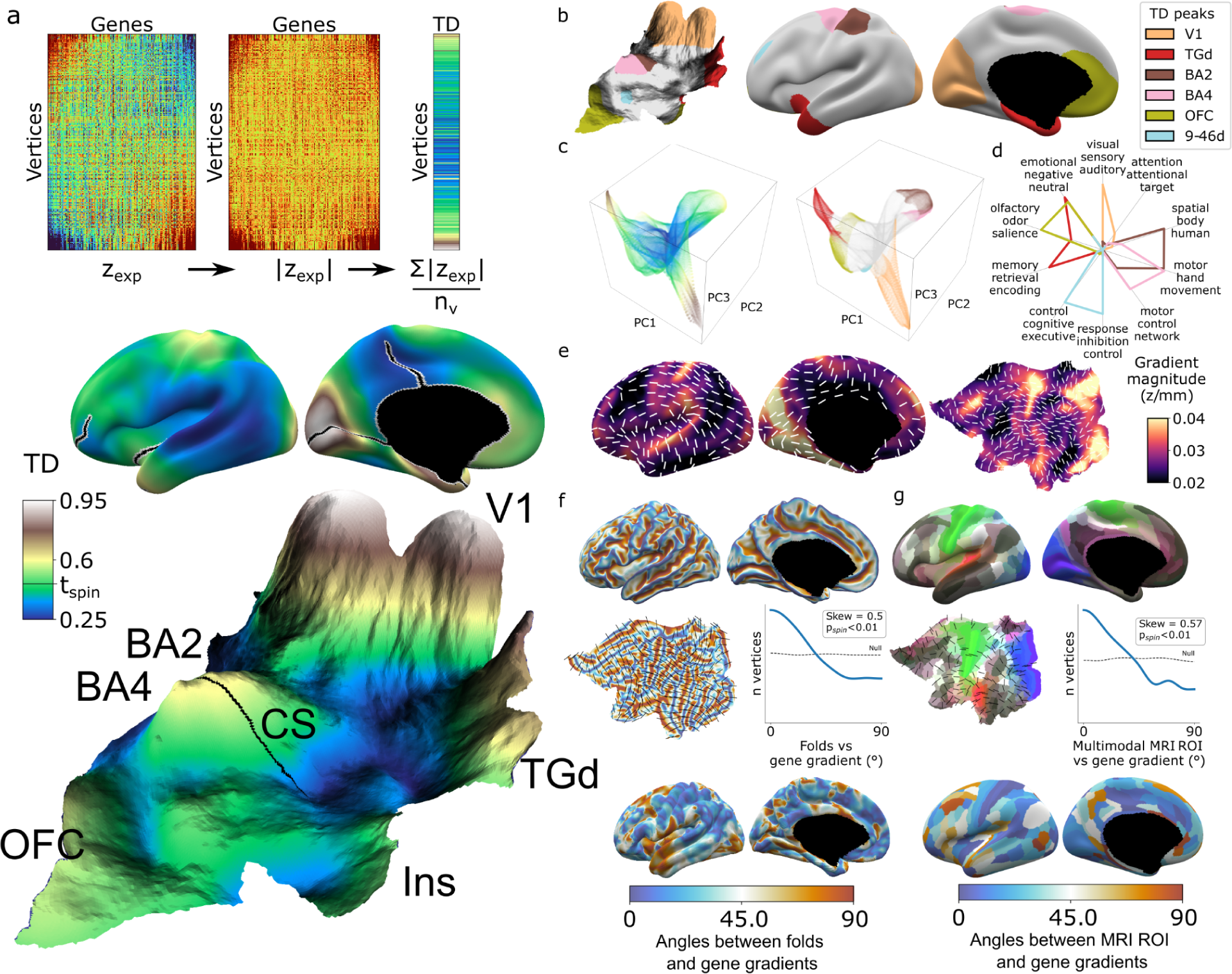
Mapping transcriptional distinctiveness in the human cortex and its alignment with macroscale structure and function. **a,** Regional transcriptomic distinctiveness (TD) can be quantified as the mean absolute z-score of dense expression map (DEM) values at each vertex (top), and visualized as a continuous cortical map (middle, TD encoded by color) or in a relief map of the flattened cortical sheet (bottom, TD encoded by color and elevation, **Sup Movie 1**). Black lines on the inflated view identify cuts for the flattening procedure. The cortical relief map is annotated to show the central sulcus (CS), and peaks of TD overlying dorsal sensory and motor cortices (Brodmann Areas, BA2, BA4), the primary visual cortex (V1), temporal pole (TGd), insula (Ins) and ventromedial prefrontal cortex (OFC). **b,** Thresholding the TD map through spatial permutation of DEMs (t_spin_ **Methods**) and clustering significant vertices by their expression profile defined six TD peaks in the adult human cortex (depicted as coloured regions on terrain and inflated cortical surfaces). **c,** Cortical vertices projected into a 3D coordinate system defined by the first 3 principal components (PCs) of gene expression, coloured by the continuous TD metric (left) and TD peaks (right). TD peaks are focal anchors of cortex-wide expression PCs **d,** TD peaks show statistically-significant functional specializations in a meta-analysis of in vivo functional MRI data. **e,** The average magnitude of local expression transitions across genes (color) and principal orientation of these transitions (white bars) varies across the cortex. **f,** Cortical folds in AHBA donors (top surface maps and middle flat-map) tend to be aligned with the principal orientation of TD change across cortical vertices (p<0.01, middle histogram, sulci running perpendicular to TD change), and the strength of this alignment varies between cortical regions. **g**, Putative cortical areas defined by a multimodal in vivo MRI parcellation of the human cortex(Glasser et al., 2016) (top surface maps and middle flat-map) also tend to be aligned with the principal direction of gene expression change across cortical vertices (p<0.01, middle histogram, sulci running perpendicular to long axis of area boundaries), and the strength of this alignment varies between cortical areas.

Integration with principal component analysis of DEMs across vertices (**Methods, Fig S3d,e**) showed that TD peaks constitute sharp poles of more recently-recognized cortical expression gradients(Burt et al., 2018) (**Fig. 2c**). The “area-like” nature of these TD peaks is reflected by the steep slopes of transcriptional change surrounding them (**Figure 2a,e**), and could be quantified as TD peaks being transcriptomically more distinctive than their physical distance from other cortical regions would predict (**Fig. S3f,g**). In contrast, transitions in gene expression are more gradual and lack such sharp transitions in the cortical regions between TD peaks (**Fig 2a,c,e, Fig S3j**). Thus, because DEMs provide spatially fine-grained estimates of cortical expression and expression change, they offer an objective framework for arbitrating between area-based and gradient-based views of cortical organization in a regionally-specific manner.

The TD peaks defined above exist as both discrete patches of cortex and the distinctive profile of gene expression which defines each peak, and this duality offers an initial bridge between macro- and microscale views of cortical organization. Specifically, we found that each TD peak overlapped with a functionally-specialized cortical region based on meta-analysis of in vivo functional neuroimaging data(Yarkoni et al., 2011) (**Methods, Fig. 2d, Table S3**), and featured a gene expression signature that was preferentially enriched for a distinct set of biological processes, cell type signatures and cellular compartments (**Methods, Table S2**). For example, the peaks overlapping area TGd and OFC were enriched for synapse-related terms, while BA2 and BA4 TD peaks were predominantly enriched for metabolic and mitochondrial terms. At a cellular level, V1 closely overlapped with DEMs for marker genes of the Ex3 neuronal subtype known to be localized to V1(Lake et al., 2016), while BA4 closely overlapped Betz cell markers(Bakken et al., 2021) (**Fig S3h**).

The expression profile of each TD peak was achieved through surrounding zones of rapid transcriptional change (**Fig 2a,e, Fig S3i,j**). We noted that these transition zones tended to overlap with cortical folds - suggesting an alignment between spatial orientations of gene expression and folding. To formally test this idea we defined the dominant orientation of gene expression change at each vertex (**Methods, Fig 2e**) and computed the angle between this and the orientation of folding (**Methods**). The observed distribution of these angles across vertices was significantly skewed relative to a null based on random alignment between angles (p_spin_<0.01, **Fig 2f, Methods**) - indicating that there is indeed a tendency for cortical sulci and the direction of fastest transcriptional change to run perpendicular to each other (p_spin_<0.01, **Fig 2f**). A similar alignment was seen when comparing gradients of transcriptional change with the spatial orientation of putative cortical areas defined by multimodal functional and structural in vivo neuroimaging(Glasser et al., 2016) (expression change running perpendicular to area long-axis, p_spin_<0.01, **Fig 2g, Methods**). Visualizing these expression-folding and expression-areal alignments revealed greatest concordance over sensorimotor, medial occipital, cingulate, and posterior perisylvian cortices (with notable exceptions of transcription change running parallel to sulci and the long-axis of putative cortical areas in lateral temporoparietal and temporopolar regions). As a preliminary probe for causality, we examined the developmental ordering of regional folding and regional transcriptional identity. Mapping the expression of high-ranking TD genes in fetal cortical laser dissection microarray data(Miller et al., 2014) from 21 PCW (Post Conception Weeks) (**Methods**) showed that the localized transcriptional identity of V1 and TGd regions in adulthood is apparent during the fetal periods that folding topology begins to emerge(Chi et al., 1977; Xu et al., 2022) (**Fig S3k**). Thus, the unique capacity of DEMs to resolve local orientations of expression change reveals a close spatial alignment between regional transitions of cortical gene expression at microscale and regional transitions of cortical folding, structure and function at macroscale.

### Cortical gene coexpression integrates diverse spatial scales of human brain organization

To complement the TD analyses above (**Fig 2**), we next used weighted gene co-expression network analysis (WGCNA(Langfelder and Horvath, 2008), **Methods**, **Fig 3a**) to achieve a more systematic integration of macro- and macroscale cortical features. Briefly, WGCNA constructs a constructs a connectivity matrix by quantifying pairwise co-expression between genes, raising the correlations to a power (here 6) to emphasize strong correlations while penalizing weaker ones, and creating a Topological Overlap Matrix (TOM) to capture both pairwise similarities expression and connectivity. Modules of highly interconnected genes are identified through hierarchical clustering. The resultant WGCNA modules enable topographic and genetic integration because they each exist as both (i) a single expression map (eigenmap) for spatial comparison with neuroimaging data (**Fig 3a,b**, **Methods**) and, (ii) a unique gene set for enrichment analysis against marker genes systematically capturing multiple scales of cortical organization, namely: cortical layers, cell types, cell compartments, protein-protein interactions (PPI) and GO terms (**Methods**, **Table S2 and S4**). Furthermore, whereas prior applications of WGCNA to AHBA data have revealed gene sets that covary in expression across many different compartments of the brain(Hartl et al., 2021; Hawrylycz et al., 2015; Kelley et al., 2018), using DEMs as input to WGCNA generates modules that are purely based on the fine-scale coordination of gene expression across the cortex. Using WGCNA, we identified 16 gene modules (M1-M16), which we then deeply annotated against independent measures of cortical organization at diverse spatial scales and developmental epochs (**Fig 3c, Methods**). Module eigenmaps were primarily driven by highly reproducible genes (**Fig S4a**) as were enrichments for annotational gene sets (median reproducibility of enriching genes=0.59, p<0.001 elevated vs. background).

**Figure 3.**
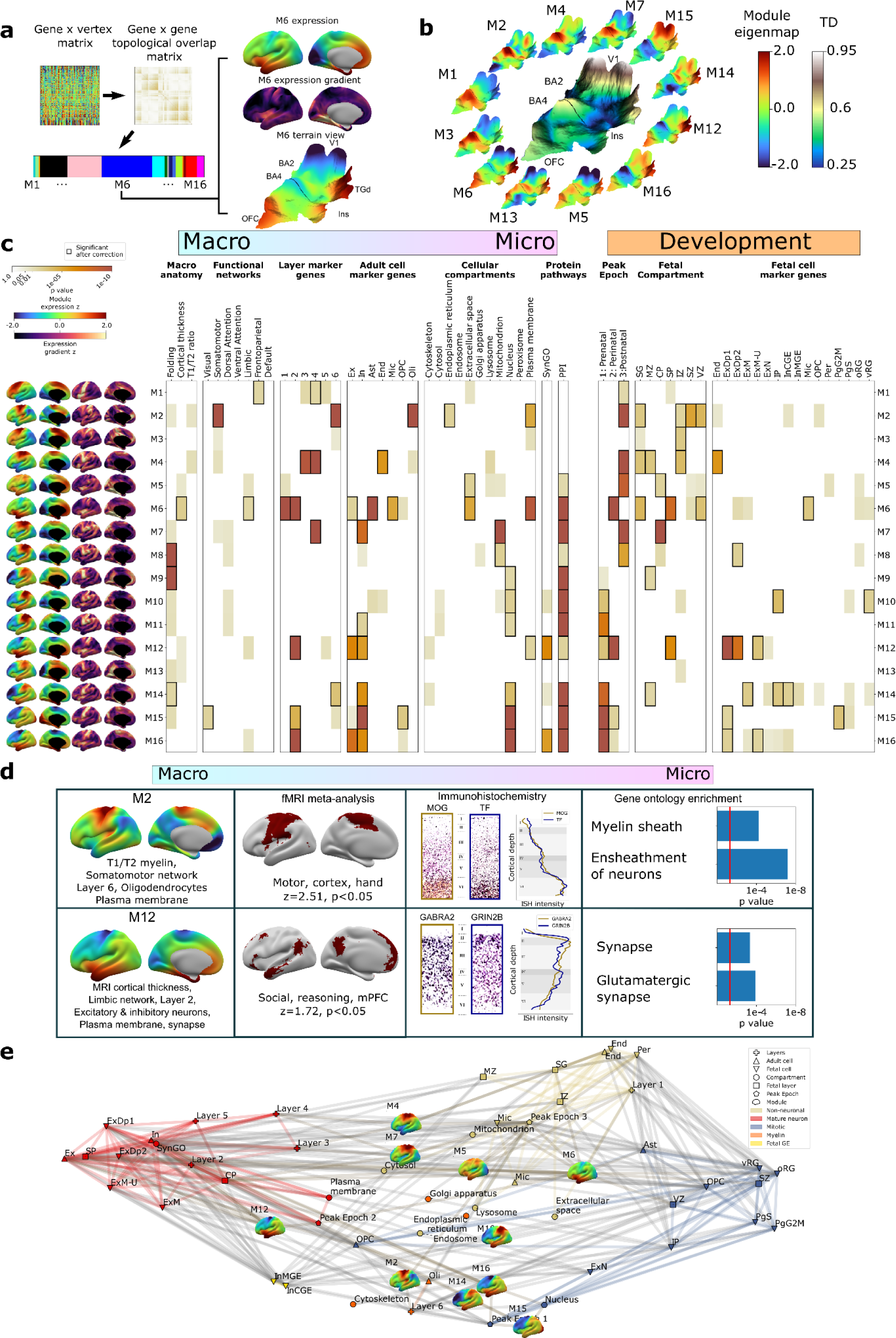
Cortex-wide Gene Coexpression Patterns Reflect Multiple Spatial Scales and Developmental Epochs of Brain Organization. **a,** Overview of Weighted Gene Co-expression Network Analysis (WGCNA) pipeline applied to the full DEM dataset. Starting top left: the pairwise DEM spatial correlation matrix is used to generate a topological overlap matrix between genes (middle top) which is then clustered. Of the 23 WGCNA-defined modules, 7 were significantly enriched for non-cortical genes and removed, leaving 16 modules. Each module is defined by a set of spatially co-expressed genes, for which the principal component of expression can be computed and mapped at each cortical point (eigenmap). M6 is shown as an example projected onto an inflated left hemisphere (M6 z-scored expression and M6 expression change), and the bulk transcriptional distinctiveness (TD) terrain view from Fig 2 (M6 expression). **b,** The extremes of WGCNA eigenmaps highlight different peaks in the cortical terrain: the main TD terrain colored by TD value (center, from Fig 2), surrounded by TD terrain projections of selected WGCNA eigenmaps. **c,** WGCNA modules (eigenmaps and gradient maps, rows) are enriched for multiscale aspects of cortical organization (columns). Cell color intensity indicates pairwise statistical significance (p<0.05), while black outlines show significance after correction for multiple comparisons across modules. Columns capture key levels of cortical organization at different spatial scales (arranged from macro-to microscale) and developmental epochs: spatial alignment between module eigenmaps and in vivo MRI maps of cortical folding orientation, cortical thickness and T1/T2 ratio, fMRI resting-state functional networks; enrichment for module gene sets for independent annotations (**Table S2**) marking: cortical layers(He et al., 2017; Maynard et al., 2021); cell types(Darmanis et al., 2015; Habib et al., 2017; Hodge et al., 2019; Lake et al., 2018, 2016; Li et al., 2018; Ruzicka et al., 2021; Velmeshev et al., 2019; Zhang et al., 2016); subcellular compartments(Binder et al., 2014); synapse-related genes(Koopmans et al., 2019); protein-protein interactions between gene products (Szklarczyk et al., 2019); temporal epochs of peak expression(Werling et al., 2020) [“fetal”: 8-24 21 post conception weeks (PCW) / “perinatal’’ 24 PCW-6 months / “postnatal” >6 months]; transient layers of the mid-fetal human cortex at 21 post conception weeks (PCW)(Miller et al., 2014)[subpial granular zone (SG), marginal zone (MZ), cortical plate (CP), subplate (SP), intermediate zone (IZ), subventricular zone (SZ) and ventricular zone (VZ)]; and fetal cell types at 17-18 PCW(Polioudakis et al., 2019). **d,** Independent validation of multiscale enrichments for selected modules M2 & M12. M2 significantly overlaps the Neurosynth topic associated with the terms motor, cortex and hand. Two high-ranking M2 genes, MOG & TF exhibit clear layer VI peaks on ISH and GO enrichment analysis myelin-related annotations. M12, overlapping the limbic network most closely overlapped the Neurosynth topic associated with social reasoning. Two high-ranking M22 genes GABRA2 and GRIN2B showed layer II ISH peaks and GO enrichment analysis revealed synaptic annotations. **e**, Network visualization of pairwise overlaps between annotational gene sets used in Fig 3c, including WGCNA module gene sets (inset expression eigenmaps).

Several WGCNA modules showed statistically significant alignments with structural and functional features of the adult cerebral cortex from in vivo imaging (**Methods**, **Fig 3c**(Glasser and Van Essen, 2011; Yeo et al., 2011)). For example, (i) the M6 eigenmap was significantly positively correlated with in vivo measures of cortical thickness from sMRI and enriched within a limbic functional connectivity network defined by resting state functional connectivity MRI, and (ii) the M8, M9 and M14 eigenmaps showed gradients of expression change that were significantly aligned with the orientation of cortical folding (especially around the central sulcus, medial prefrontal and temporo-parietal cortices, **Fig S4b**). At microscale, several WGCNA module gene sets showed statistically significant enrichments for genes marking specific cortical layers(He et al., 2017; Maynard et al., 2021) and cell types(Darmanis et al., 2015; Habib et al., 2017; Hodge et al., 2019; Lake et al., 2018, 2016; Li et al., 2018; Ruzicka et al., 2021; Velmeshev et al., 2019; Zhang et al., 2016) (**Methods**, **Fig 3c, Table S4**). These microscale enrichments were often congruent between cortical layers and cell classes annotations, and in keeping with the linked eigenmap (**Fig 3c**, **Table S4**). For example, M4 - which was uniquely co-enriched for markers of endothelial cells and middle cortical layers - showed peak expression over dorsal motor cortices which are known to show expanded middle layers(Bakken et al., 2021; Wagstyl et al., 2020) with rich vascularization(Pfeifer, 1940) relative to other cortical regions. Similarly, M6 - which was enriched for markers of astrocytes, microglia and excitatory neurons as well as layers 1/2 - showed peak expression over rostral frontal and temporal cortices which are known to possess relatively expanded supragranular layers(Wagstyl et al., 2020) that predominantly contain the apical dendrites of excitatory neurons and supporting glial cells(von Economo and Koskinas, 1925). We also observed that modules with similar eigenmaps (**Fig S4c**), (including overlaps of multiple modules with the same TD peak) could show contrasting gene set enrichments. For example M2 and M4 both showed peak expression of dorsal sensorimotor cortex (i.e. TD areas BA2 and BA4), but M2 captures a distinct architectonic signature of sensorimotor cortex from the mid-layer vascular signal of M4: expanded and heavily myelinated layer 6(Bakken et al., 2021; Palomero-Gallagher and Zilles, 2019; Wagstyl et al., 2020) (**Fig 3c**). The spatially co-expressed gene modules detected by WGCNA were not only congruently co-enriched for cortical layer and cell markers, but also for nanoscale features such as sub-cellular compartments(Binder et al., 2014) (**Table S2 and S4**) (often aligning with the cellular enrichments) and protein-protein interactions(Szklarczyk et al., 2019) (PPI) (**Methods**, **Fig 3c, Table S4**). This demonstrates the capacity of our resource to tease apart subtle subcomponents of neurobiology based on cortex-wide expression patterns.

To further assess the robustness of these multiscale relationships, we focused on two modules with contrasting multiscale signatures - M2 and M12 - and tested for reproducibility of our primary findings (**Fig 3c**) using orthogonal methods. Our primary analyses indicated that M2 has an expression eigenmap which overlaps with the canonical somato-motor network from resting-state functional neuroimaging(Yeo et al., 2011), and contains genes that are preferentially expressed in cortical layer 6 from layer-resolved transcriptomics(He et al., 2017; Maynard et al., 2021), and in oligodendrocytes from snRNAseq(Darmanis et al., 2015; Habib et al., 2017; Hodge et al., 2019; Lake et al., 2018, 2016; Li et al., 2018; Ruzicka et al., 2021; Velmeshev et al., 2019; Zhang et al., 2016) (**Fig 3c**). We were able to verify each of these observations through independent validations including: spatial overlap of M2 expression with meta analytic functional activations relating to motor tasks(Yarkoni et al., 2011); immunohistochemistry localization of high-ranking M2 genes to deep cortical layers(Zeng et al., 2012) (**Methods**); and significant enrichment of M2 genes for myelin-related GO terms (**Fig 3d, Table S4**). By contrast, our primary analyses indicated that M12 - which had peak expression over ventral frontal and temporal limbic cortices - was enriched for marker genes for layer 2, neurons and the synapse (**Fig 3c**). These multiscale enrichments were all supported by independent validation analyses, which showed that: the M12 eigenmaps is enriched in a limbic network that is activated during social reasoning(Yarkoni et al., 2011); high-ranking M12 marker genes show elevated expression in upper cortical layers by immunohistochemistry(Zeng et al., 2012) (**Methods**); and, there is a statistically-significant over representation of synapse compartment GO terms in the M12 gene set (**Fig 3d, Table S4**).

### Linking spatial and developmental aspects of cortical organization

Given that adult cortical organization is a product of development, we next asked if eigenmaps of adult cortical gene expression (**Fig 3a,b**) are related to the patterning of gene expression between fetal stages and adulthood. To achieve this, we tested WGCNA module gene sets for enrichment of developmental marker genes from 3 independent postmortem studies (rightmost columns, **Fig 3c**) capturing genes with differential expression between (i) 3 developmental epochs between 8 post-conception weeks (PCWs) and adulthood (*BrainVar* dataset from prefrontal cortex(Werling et al., 2020)) (ii) 7 histologically-defined zones of mid-fetal (21 PCW) cortex(Miller et al., 2014) (**Methods**, **Table S1 and S2),** and (iii) 16 mid-fetal (17-18 PCW) cell-types(Polioudakis et al., 2019) (**Methods, Table S2**).

Comparison with the *BrainVar* dataset revealed that most module eigenmaps (13 of all 16 cortical modules) were enriched for genes with dynamic, developmentally-coordinated expression levels between early fetal and late adult stages (**Figure 3c, Table S4**). This finding was reinforced by supplementary analyses modeling developmental trajectories of eigenmap gene set expression between 12 PCW and 40 years in the *BrainSpan* dataset(Li et al., 2018) (**Methods**, **Fig S4d**), and further qualified by the observation that several WGCNA modules were also differentially enriched for markers of mid-fetal cortical layers and cell-types(Miller et al., 2014; Polioudakis et al., 2019) (**Figure 3c, Table S4**). As observed for multiscale spatial enrichments (**Fig 3c,d**); the developmental enrichments of modules were often closely coordinated with one another, and eigenmaps with similar patterns of regional expression could possess different signatures of developmental enrichment. For example, the M6 and M12 eigenmaps shared a similar spatial expression pattern in the adult cortex (peak expression in medial prefrontal, anterior insula and medio-ventral temporal pole), but captured different aspects of human brain development that aligned with the cyto-laminar enrichments of M6 and M12 in adulthood. The M6 gene set - which was enriched for predominantly glial elements of layers 1 and 2 in adult cortex - was also enriched for markers of mid-fetal microglia(Polioudakis et al., 2019), the transient fetal layers that are known to be particularly rich in mid-fetal microglia (subpial granular, subplate, and ventricular zone(Monier et al., 2007)), and the mid-late fetal epoch when most microglial colonization of the cortex is thought to be achieved(Menassa and Gomez-Nicola, 2018) (**Fig 3c**). In contrast, the M12 gene set - which was enriched for predominantly neuronal elements of layer 2 in adult cortex - also showed enrichment for marker genes of developing fetal excitatory neurons, the fetal cortical subplate, and windows of mid-late fetal development when developing neurons are known to be migrating into a maximally expanded subplate(Molnár et al., 2019).

The striking co-enrichment of WGCNA modules for features of both the fetal and adult cortex (**Fig 3c**) implied a patterned sharing of marker genes between cyto-laminar features of the adult and fetal cortex. To more directly test this idea, and characterize potential biological themes reflected by these shared marker genes, we carried out pairwise enrichment analyses between all annotational gene sets from **Fig 3c**. These gene sets collectively draw from a diverse array of study designs encompassing bulk, laminar, and single cell transcriptomics of the human cortex between 10 PCW and 60 years of life (**Methods**(Darmanis et al., 2015; Habib et al., 2017; He et al., 2017; Li et al., 2018; Maynard et al., 2021; Miller et al., 2014; Polioudakis et al., 2019; Ruzicka et al., 2021; Velmeshev et al., 2019; Werling et al., 2020; Zhang et al., 2016)). Network visualization and clustering of the resulting adjacency matrix (**Fig S4e**) revealed an integrated annotational space defined by five coherent clusters (**Fig 3e**). A mature neuron cluster encompassed markers of post-mitotic neurons and the compartments that house them in both fetal and adult cortex (red, **Fig 3e, Table S2**, example core genes: NRXN1, SYT1, CACNG8). This cluster also included genes with peak expression between late fetal and early postnatal life, and those localizing to the plasma membrane and synapse. A small neighboring fetal ganglionic eminence cluster (Fetal GE, yellow, **Fig 3e, Table S2**, example core genes: NPAS3, DSX, DCLK2) contained marker sets for migrating inhibitory neurons from the medial and caudal ganglionic eminence in mid-fetal life. These two neuronal clusters - mature neuron and Fetal GE - were most strongly connected to the M12 gene set (**Methods**), which highlights medial prefrontal, and temporal cortices possessing a high ratio of neuropil:neuronal cell bodies(Collins et al., 2010; Spocter et al., 2012). A mitotic annotational cluster (blue, **Fig 3e, Table S2**, example core genes: CCND2, MEIS2, PHLDA1) was most distant from these two neuronal clusters, and included genes showing highest expression in early development as well as markers of cycling progenitor cells, radial glia, oligodendrocyte precursors, germinal zones of the fetal cortex, and the nucleus. This cluster was most strongly connected to the M15 gene set, which shows high expression over occipito-parietal cortices distinguished by a high cellular density and notably low expression in lateral prefrontal cortices, which possess low cellular density(Collins et al., 2016). The mature neuron and mitotic clusters were separated by two remaining annotational clusters for non-neuronal cell types and associated cortical layers. A myelin cluster (orange, **Fig 3e, Table S2**, example core genes: MOBP, CNP, ACER3) - which contained gene sets marking adult layer 6, oligodendrocytes, and organelles supporting the distinctive biochemistry and morphology of oligodendrocytes (the golgi apparatus, endoplasmic reticulum and cytoskeleton) - was most connected to the M2 gene set highlighting heavily myelinated motor cortex(Nieuwenhuys and Broere, 2017). A non-neuronal cluster (yellow, **Fig 3e, Table S2**, example core genes: TGFBR2, GMFG, A2M) - which encompassed marker sets for microglia, astrocytes, endothelial cells, pericytes, and markers of superficial adult and fetal cortical layers that are relatively depleted of neurons - was most connected to the M6 gene set highlighting medial temporal and anterior cingulate cortices with notably high non-neuronal content(Collins et al., 2010).

These analyses show that the regional patterning of bulk gene expression captures the organization of the human cortex across multiple spatial scales and developmental stages such that (i) the summary expression maps of spatially co-expressed gene sets align with independent in vivo maps of macroscale structure and function from neuroimaging, while (ii) the spatially co-expressed gene sets defining these maps show congruent enrichments for specific adult cortical layers and cell-types as well as developmental precursors of these features spanning back to mid-fetal life.

### ASD risk genes follow two different spatial patterns of cortical expression, which capture distinct aspects of cortical organization and differentially predict cortical changes in ASD

The findings above establish that gene co-expression modules in the human cortex capture multiple levels of biological organization ranging from subcellular organelles, to cell types, cortical layers and macroscale patterns of brain structure and function. Given that genetic risks for atypical brain development presumably play out through such levels of biological organization, we hypothesized that disease associated risk genes would be enriched within WGCNA module gene sets. Testing this hypothesis simultaneously offers a means of further validating our analytic framework, while also potentially advancing understanding of disease biology. To test for disease gene enrichment in WGCNA modules, we compiled lists of genes enriched for deleterious rare variants in autism spectrum disorder(Ruzzo et al., 2019; Satterstrom et al., 2020) (ASD), schizophrenia(Singh et al., 2020) (Scz), severe developmental disorders (DDD)(Deciphering Developmental Disorders Study, 2017) and epilepsy(Heyne et al., 2018) (**Table S2**). We considered rare (as opposed to common) genetic variants to focus on high effect-size genetic associations and avoid ongoing uncertainties regarding the mapping of common variants to genes(Tam et al., 2019). We observed that disease associated gene sets were significantly enriched in several WGCNA modules (**Fig 4a**), with two modules showing enrichments for more than one disease: M15 (ASD, Scz and DDD) and M12 (ASD and Epilepsy).

**Figure 4.**
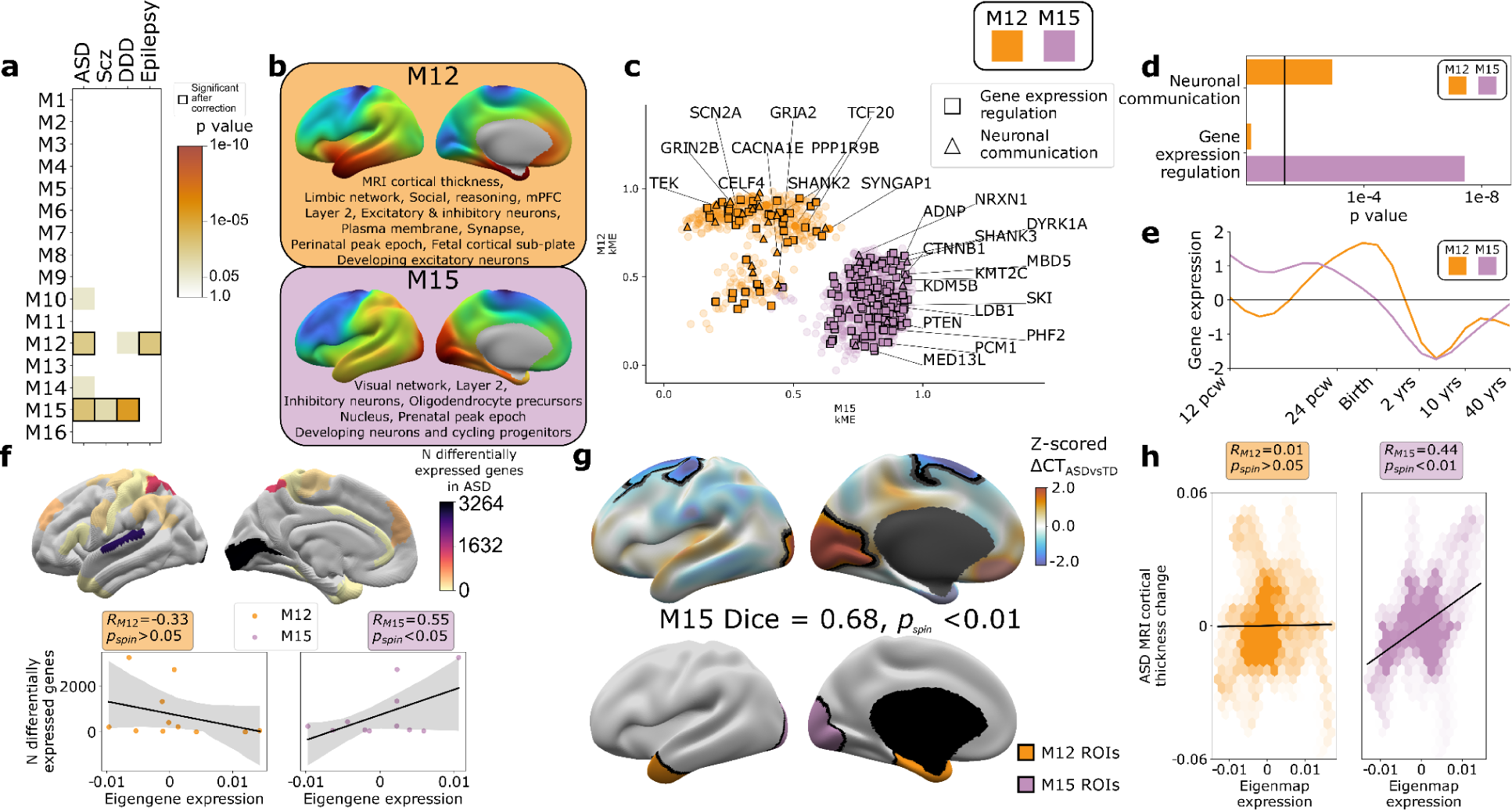
ASD risk genes follow two different spatial patterns of cortical gene expression which differentially predict cortical changes in ASD. **a,** Enrichment of WGCNA module gene sets for risk genes associated with atypical brain development through enrichment of rare deleterious variants in studies of Autism Spectrum Disorder (ASD), Schizophrenia (Scz), severe developmental disorders (DDD, Deciphering Developmental Disorders study) and Epilepsy. Cell color intensity indicates pairwise statistical significance (p<0.05), while outlined matrix cells survived correction for multiple comparisons across modules. **b**, Summary of multiscale and developmental annotations from Fig 3c for M12 and M15: the only two WGCNA modules enriched for risk genes of more than one neurodevelopmental disorder. **c**, M12 and M15 genes clustered by the strength of their membership to each module. Color encodes module membership. Shape encodes annotations for two GO Biological Process annotations that differ between the module gene sets: neuronal communication and regulation of gene expression. Text denotes specific ASD risk genes. **d**, contrasting GO enrichment of M12 and M15 for neuronal communication and regulation of gene expression GO Biological Process annotations. **e**, M12 and M15 differ in the developmental trajectory of their average cortical expression between early fetal and mid-adult life(Li et al., 2018). **f**, Regional differences in intrinsic expression of the M15 module (but not the M12 module) in adult cortex is correlated with regional variation in the severity of altered cortical gene expression (number of differentially expressed genes) in ASD(Haney et al., 2020). **g**, Statistically-significant regional alterations of cortical thickness (CT) in ASD compared to typically developing controls from in vivo neuroimaging(Di Martino et al., 2017, 2013) (top). Areas of cortical thickening show a statistically-significant spatial overlap (Dice overlap = 0.68, p_spin_<0.01) with regions of peak intrinsic expression for M15 in adult cortex (bottom). **h**, M15 eigenmap expression (but not M12 eigenmap) shows significant spatial correlation with relative cortical thickness change in ASD.

ASD was the only disorder to show a statistically-significant enrichment of risk genes within both M12 and M15 (**Fig 4a**) - providing an ideal setting to ask if and how this partitioning of ASD risk genes maps onto (i) multiscale brain organization in health, and (ii) altered brain organization in ASD. The eigenmaps and gene set enrichments of M12 vs. M15 implicated two contrasting multiscale motifs in the biology of ASD (**Fig 4b**). ASD risk genes including SCN2A, SYNGAP1, and SHANK2 resided within the M12 module (**Fig 4c**) which is most highly expressed within a distributed cortical system that is activated during social reasoning tasks (p_spin_<0.01, **Fig 3c,d**, **Fig 5b**). The M12 gene set is also enriched for: genes with peak cortical expression in late-fetal and early postnatal life; marker genes for the fetal subplate and developing excitatory neurons; markers of layer 2 and mature neurons in adult cortex; and synaptic genes involved in neuronal communication (**Fig 3c,d**, **Fig 4b,c,d,e**, **Table S4**). In contrast, ASD risk genes including ADNP, KMT5B, and MED13L resided within the M15 module (**Fig 4c**), which is most highly expressed in primary visual cortex and associated ventral temporal pathways for object recognition/interpretation(Kravitz et al., 2013) (p_spin_<0.05, **Fig 3c,d**, **Fig 4b, Table S4**). The M15 module is also enriched for: genes showing peak cortical expression in early fetal development, marker genes for cycling progenitor cells in the fetal cortex; markers of layer 2, inhibitory neurons and oligodendrocyte precursors in the adult cortex (**Fig 3c,d**, **Fig 4b,c,d,e, Table S4**). The alignment of ASD risk genes with M12 and M15 was reinforced when considering all 135 ASD risk genes: spatial co-expression analyses split these genes into two clear subsets with mean expression maps that most closely resembled M12 & M15 (**Fig S5a,b**). Thus - using only spatial patterns of cortical gene expression in adulthood, our analytic framework was able to recover the previous PPI and GO-based partitioning of ASD risk genes into synaptic vs. nuclear chromatin remodeling pathways(Parikshak et al., 2013; Satterstrom et al., 2020), and then place these pathways into a richer biological context based on the known multiscale associations of M12 and M15 (**Figs 3c, 4a**).

We next sought to address whether regional differences in M12 and M15 expression were related to regional cortical changes observed in ASD. To test this idea, we used two orthogonal indices of cortical change in ASD that capture different levels of biological analysis - the number of differentially expressed genes (DEGs) postmortem(Haney et al., 2020), and the magnitude of changes in cortical thickness (CT) as measured by in vivo sMRI(Di Martino et al., 2017). Regional DEG counts were derived from a recent postmortem study of 725 cortical samples from 11 cortical regions in 112 ASD cases and controls(Haney et al., 2020), and compared with mean M12 and M15 expression within matching areas of a multimodal MRI cortical parcellation(Glasser et al., 2016). The magnitude of regional transcriptomic disruption in ASD was statistically-significantly positively correlated with region expression of the M15 module (r=0.6, p_spin_<0.05), but not the M12 module (r=-0.3, p_spin_>0.05) (**Fig 4f**). This dissociation is notable because M15 (but not M12) is enriched for genes involved in the regulation of gene expression (**Fig 4d**). Thus the enrichment of regulatory ASD risk genes within M15, and the intrinsically high expression of M15 in occipital cortex may explain why the occipital cortex is a hotspot of altered gene expression in ASD.

To compare M12 and M15 expression with regional variation in cortical anatomy changes in ASD, we harnessed the multicenter ABIDE datasets containing brain sMRI scans from 751 participants with idiopathic ASD and 773 controls(Di Martino et al., 2017, 2013). We preprocessed all scans using well-validated tools for harmonized estimation of cortical thickness (CT)(Fischl, 2012) from multicenter data (**Methods**), and then modeled CT differences between ASD and control cohorts at 150,000 points (vertices) across the cortex (**Methods**). This procedure revealed two clusters of statistically-significant CT change in ASD (**Methods**, **Fig 4g**, upper panel) encompassing visual and parietal cortices (relative cortical thickening vs. controls) as well as superior frontal vertices (relative cortical thinning). The occipital cluster of cortical thickening in ASD showed a statistically-significant spatial overlap with the cluster of peak M15 expression (**Fig 4g**, upper panel, **Methods**, Dice coefficient = 0.7, p_spin_<0.01), and relative cortical thickness change correlated with the M15 eigenmap (**Fig 4h**). In contrast, M12 expression was not significantly aligned with CT change in ASD (**Fig 4g,h**). Testing these relationships in the opposite direction - i.e. asking if regions of peak M12 and M15 expression are enriched for directional CT change in ASD relative to other cortical regions - recovered the M15-specific association with regional cortical thickening in ASD (**Fig S5c**).

Taken together, the above findings reveal that an occipital hotspot of altered gene expression and cortical thickening in ASD overlaps with an occipital hotspot of high expression for a subset of ASD risk genes. These ASD risk genes are spatially co-expressed in a module enriched for several connected layers of biological organization (**Fig 3c, 4b,c,d**) spanning: nuclear pathways for chromatin modeling and regulation of gene expression; G2/M phase cycling progenitors and excitatory neurons in the mid-fetal cortex; oligodendrocytes and layer 2 cortical neurons in adult cortex; and occipital functional networks involved in visual processing. These multiscale aspects of cortical organization can now be prioritized as potential targets for a subset of genetic risk factors in ASD, and the logic of this analysis in ASD can now be generalized to any disease genes of interest.

## Discussion

We build on the most anatomically comprehensive dataset of human cortex gene expression available to date(Hawrylycz et al., 2012), to generate, validate, characterize, apply and share spatially dense measures of gene expression that capture the topographically continuous nature of the cortical mantle. By representing patterns of human cortical gene expression without the imposition of a priori boundaries(Burt et al., 2018; Hawrylycz et al., 2015) our library of dense gene expression maps (DEMs) allows anatomically-unbiased analyses of local gene expression levels as well as the magnitudes and directions of local gene expression change. This core spatial property of DEMs unlocks several methodological and biological advances. First, the unparcellated nature of DEMs allows us to agnostically define cortical zones with extreme transcriptional profiles or unusually rapid transcriptional change - which we show to capture microstructural cortical properties and align with folding and functional specializations at the macroscale (**Fig 2**). By establishing that some of these cortical zones are evident at the time of cortical folding, we lend support to a “protomap”(O’Leary, 1989; O’Leary et al., 2007; Rakic, 1988; Rakic et al., 2009) like model where the placement of some cortical folds is set-up by rapid tangential changes in cyto-laminar composition of the developing cortex(Ronan et al., 2014; Toro and Burnod, 2005; Van Essen, 2020). The DEMs are derived from fully folded adult donors, and therefore some of the measured genetic-folding alignment might also be induced by mechanical distortion of the tissue during folding(Heuer and Toro, 2019; Llinares-Benadero and Borrell, 2019). However, no data currently exist to conclusively assess the directionality of this gene-folding relationship.

We show that DEMs can recover sharp boundaries in gene expression despite being generated by interpolation algorithms that do not explicitly encode step-changes in expression between cortical regions. This property of DEMs will help to target future studies of human cortical patterning (for example directing single cell and spatial omics resources), and we illustrate this utility by applying DEMs to discover two new expression boundaries in the human cortex. Second, we use spatial correlations between DEMs to decompose the complex topography of cortical gene expression into a smaller set of cortex-wide transcriptional programs that capture distinct aspects of cortical biology - at multiple spatial scales and multiple developmental epochs (**Fig 3**). This effort provides an integrative model that links expression signatures of cell-types and layers in prenatal life to the large-scale patterning of regional gene expression in the adult cortex, which can in turn - through DEMs - be compared to the full panoply of in vivo brain phenotypes provided by modern neuroimaging. Indeed, future work might find direct links between these module eigenvectors and similar low-frequency eigenvectors of cortical geometry have been used as basis functions to segment the cortex (Lefèvre et al., 2018) and explain complex functional activation patterns(Pang et al., 2023). Third, we find that some of these cortex-wide expression programs in adulthood are enriched for disease risk genes - which offers a new path to nominating candidate disease mechanisms across different levels of biological organization (**Fig 4**). For example the M15 module defines a normative spatial pattern of cortical gene co-expression which not only captures a functionally-enriched subset of ASD genes(Satterstrom et al., 2020), but also shows multiscale enrichments and regionally-specific expression patterns that tie together several independently-reported aspects of ASD neurobiology. Specifically, M15 newly integrates (i) the concentration of ASD risk genes and dysregulated gene expression in upper-layer excitatory neurons(Velmeshev et al., 2019), (ii) the accentuation of altered gene expression and thickness in occipital cortical regions, and (iii) the early emergence amongst children at heightened genetic risk for ASD of behaviorally-relevant changes in cortical structure and function(Girault et al., 2022) within occipital systems important for the processing of visual information. Crucially, the strategy applied in our analysis of ASD risk genes can be generalized to risk genes for any brain disorder of interest to place known risk factors for disease into the rich context of multiscale cortical biology.

Finally, the collection of DEMs, annotational gene sets and statistical tools used in this work is shared as a new resource to accelerate multiscale neuroscience by allowing flexible and spatially unbiased translation between genomic and neuroanatomical spaces. Of note, this resource can easily incorporate any future expansions of brain data in either neuroanatomical or genomic space. We anticipate that it will be particularly valuable to incorporate new data from the nascent, but rapidly expanding fields of high throughput histology(Wagstyl et al., 2020), single cell-omics(Bakken et al., 2021), and large-scale imaging-genetics studies(Smith et al., 2021). Taken together, MAGICC enables a new integrative capacity in the way we study the brain, and hopefully serves to spark new connections between previously distant datasets, ideas and researchers.

## Materials and Methods

### Materials and Methods overview

1. **Creating spatially dense maps of human cortical gene expression (Fig 1a-d)**
2. **Benchmarking dense expression maps (DEMs)**

a. Spin tests for comparing two spatial maps
b. Replicability and independence from cortical sampling density (Fig S1).
c. Alignment with reference measures of cortical organization (Fig 1 e-g)
3. **Characterizing the topography of DEMs**

a. Transcriptomic distinctiveness (TD) and principal component analysis (Fig 2a-c)
b. Relating adult TD peaks to fetal gene expression (Fig S3j)
c. Local gradient analysis (Fig 2e-g)
d. Weighted Gene Co-expression Network Analysis (WGCNA) (Fig 3a-c)
4. **Multiscale annotation of WGCNA modules (Fig 3c,d)**

a. Map-based annotations
b. Gene-set based annotations
5. **Combining gene-set based annotations of the cortical sheet (Fig 3e, Fig S3d)**
6. **Disease enrichment and ASD-based analysis of WGCNA modules**

a. Characterizing ASD gene enrichments in M12 and M15
b. Comparing M12 and M15 expression to regional changes of cortical gene expression in ASD (Fig 4f)
c. Comparing M12 and M15 expression to regional changes of cortical thickness in ASD (Fig 4g, h, Fig S5c)
7. **Preprocessing and analysis of structural MRI data**

a. AHBA donors
b. OASIS (Fig 1e)
c. ABIDE

### 1. Creating spatially dense maps of human cortical gene expression (Fig 1a-d)

Cortical surfaces were reconstructed for each AHBA donor MRI using FreeSurfer(Fischl, 2012), and coregistered between donors using surface matching of individuals’ folding morphology (MSMSulc) (Robinson et al., 2018). An average donor cortical mesh was also created for analyses of cortical morphology, by averaging the vertex coordinates of volumetrically aligned meshes for the 6 donors.

Probe-level data measures of gene expression for all samples in the AHBA adult brain microarray dataset were downloaded from (https://human.brain-map.org/static/download) - providing log2-transformed measures of gene expression for 58,692 probes in each of 3,702 brain tissue samples from six donors (**Table S1**). Within- and across-brain normalization of these probe level gene expression values was implemented as detailed by the Allen Institute for Brain Science White Paper (http://help.brain-map.org/download/attachments/2818165/WholeBrainMicroarray_WhitePaper.pdf). Probes were reannotated using the updated manifest from Arnautkevic et al(Arnatkeviciute et al., 2019), excluding genes lacking an Entrez, and probe-level expression values were averaged for each gene to yield a single gene*sample expression matrix for each donor. As only 2 donors had measurements from right hemispheres, samples were filtered by region to retain those originating from the cerebral cortex left hemisphere only. This decision was made given evidence for potential asymmetries in gene expression within the human cortex(de Kovel et al., 2018), and known differences in cortical shape between the hemispheres that complicate the reflection of sample locations from left to right cortical sheets(Jo et al., 2012). The above steps resulted in a final set of 6 donor-level gene*sample matrices from the left cerebral cortex for downstream analyses. These matrices collectively contained scaled expression values for 20,781 genes in each of 1304 cortical samples.

Native subject MRI coordinates were extracted for every cortical sample in each donor (**Fig 1a**). Nearest mid-surface cortical vertices were identified for each sample, excluding samples further than 20mm from a cortical coordinate. For cortical vertices with no directly sampled expression, expression values were interpolated from their nearest sampled neighbor vertex on the spherical surface (Moresi and Mather, 2019) (**Fig 1b**). Sampling density ρ in each subject was calculated as the number of samples per mm^2^, from which average inter-sample distance, *d,* was estimated using the formula: 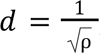, giving a mean intersample distance of 17.7mm ± 1.2mm. Surface expression maps were smoothed using the Connectome Workbench toolbox (Glasser et al., 2013) with a 20mm full-width at half maximum Gaussian kernel, selected to be consistent with this sampling density (**Fig 1c**). To align subjects’ expression, expression values were z-scored by the mean and standard deviation across vertices (given the known criticality of within-subject scaling of AHBA expression values (Markello et al., 2021)) and then averaged across the 6 subjects (**Fig 1d**) - yielding spatially dense estimates of expression at 29696 vertices across the left cerebral cortex per gene. Vertex-wise, rather than sample-level, estimation of mean and standard deviation mitigates potential biases introduced by intersubject variability in the regional distribution and density of cortical samples. For Y-linked genes, DEMs were calculated from male donors only. For each of the resulting 20,781 gene-level expression maps, the orientation and magnitude of gene expression change at each vertex (i.e. the gradient) was calculated for folded, inflated, spherical and flattened mesh representations of the cortical sheet using Connectome Workbench’s metric gradient command (Glasser et al., 2013).

## 2. Benchmarking dense expression maps (DEMs)

### a. Spin tests for comparing two spatial maps

Cortical maps exhibit spatial autocorrelation that can inflate the False Positive Rate, for which a number of methods have been proposed(Alexander-Bloch et al., 2018; Burt et al., 2020; Vos de Wael et al., 2020). At higher degrees of spatial smoothness, this high False Positive Rate is most effectively mitigated using the spin test(Alexander-Bloch et al., 2018; Markello and Misic, 2021; Vos de Wael et al., 2020). In the following analyses when generating a test statistic comparing two spatial maps, to generate a null distribution, we computed 1000 independent spins of the cortical surface using https://netneurotools.readthedocs.io, and applied it to the first map whilst keeping the second map unchanged. The test statistic was then recomputed 1000 times to generate a null distribution for values one might observe by chance if the maps shared no common organizational features. This is referred to throughout as the “spin test” and the derived p-values as p_spin_.

An additional null dataset was generated to test whether intrinsic geometry of the cortical mesh and its impact on interpolation for benchmarking analyses of DEMs and gradients (Fig S1d, Fig S2d, Fig S3c). In these analyses, the original samples were rotated on the spherical surface prior to subsequent interpolation, smoothing and gradient calculation. Due to computational constraints the full dataset was recreated only for 10 independent spins. These are referred to as the “spun+interpolated null”.

### b. Replicability and independence from cortical sampling density (Fig S1)

We assessed the replicability of DEMs by applying the above steps for DEM generation to non-overlapping donor subsets and comparing DEMs between the resulting sub-atlases. We quantified DEM agreement between sub-atlases at both the gene-level (correlation in expression across vertices for each gene, **Fig S1c**) and the vertex-level (correlation in ranking of genes by their scaled expression values at each vertex, **Fig S1d,e**). These sub-atlas comparisons were done between all possible pairs of individuals, donor duos and donor triplets to give distributions and point estimates of reproducibility for atlases formed of 1, 2 and 3 donors. Learning curves were fitted to these data to estimate the projected gene-level and vertex-level DEM reproducibility of our full 6-subject sample atlas(Figueroa et al., 2012)(**Fig S1c**).

To assess the effect of data interpolation in DEM generation we compared gene-level and vertex-level reproducibility of DEMs against a “ground truth” estimate of these reproducibility metrics based on uninterpolated expression data. To achieve a strict comparison of gene expression values between different individuals at identical spatial locations we focused these analyses on the subset of AHBA samples where a sample from one subject was within 3 mm geodesic distance of another. This resulted in 1097 instances (spatial locations) with measures of raw gene expression of one donor, and predicted values from the second donor’s un-interpolated AHBA expression data and interpolated DEM. We computed gene-level and vertex-level reproducibility of expression using the paired donor data at each of these sample points - for both DEM and uninterpolated AHBA expression values. By comparing DEM reproducibility estimates with those for uninterpolated AHBA expression data, we were able to quantify the combined effect of interpolation and smoothing steps in DEM generation. We used gene-level reproducibility values from DEMs and uninterpolated AHBA expression data to compute a gene-level difference in reproducibility, and we then visualized the distribution of these difference values across genes (**Fig S1a**). We used gene-rank correlation to compare vertex-level reproducibility values between DEMs and uninterpolated AHBA expression data (**Fig S1b**).

Theoretically, regional gradients of expression change in DEMs could be biased by regional variations in the density of AHBA cortical sampling. To test for this, in each individual subject, we calculated the spatial relationship between the sampling density and mean gene gradient magnitude (**Fig S1g**). We additionally tested whether the regional variability of gene rank predictability in the atlas (shown in **Fig S1f**) was linked to the sampling density within the atlas.

### c. Alignment with reference measures of cortical organization (Fig 1 e-g)

We first determined if our DEM library was able to differentiate between genes that are known to show cortical expression (CExp) and those without any prior evidence of cortical expression (NCExp) - motivated by the strong expectation that NCExp genes should lack a consistent spatial gradient in expression. For this test, we defined a set of 16573 CExp genes by concatenating the genes coding for proteins found in the “cortex” tissue class of the human protein atlas(Sjöstedt et al., 2020) genes identified as markers for cortical layers or cortical cells (see below,(Darmanis et al., 2015; Habib et al., 2017; He et al., 2017; Hodge et al., 2019; Lake et al., 2018, 2016; Li et al., 2018; Maynard et al., 2021; Ruzicka et al., 2021; Velmeshev et al., 2019; Zhang et al., 2016)). The remaining 4,208 genes in our DEM library were classified as NCExp. Fisher’s exact test was used to assess whether genes with lower gene reproducibility (r<0.5) were enriched for NCExp genes. We projected vertex-level reproducibility values for CExp and NCExp genes onto the cortical surface for visual comparison, and also computed the mean cross-vertex reproducibility for each of these maps (**Fig S1f**).

We next compiled data from independent studies for a range of macroscale and microscale cortical features that would be expected to align with specific DEM maps, and asked if the spatial patterns of cortical gene expression from DEMs showed the expected alignment with these independent data. These independent comparison studies were selected to span diverse measurement methods and data modalities representing a range of spatial scales.

We first sought to establish whether local changes in DEMs, i.e. the gradient maps of gene expression, could be used to validate existing areal border genes and identify novel candidates. Using a parcellation of the cortex based on multimodal structural and functional neuroimaging (Glasser et al., 2016), we identified the vertices along the boundary between a pair of regions (e.g. V1 & V2). The mean DEM gradient at these vertices was quantified for each gene, enabling us to rank genes by their exhibited border-like features at this cortical location. We then assessed the ranking of known lists of areal marker genes for a given border against a randomly sampled null distribution. To validate known areal marker genes derived from previous ISH studies, we took examples from the human visual cortex (Zeng et al., 2012), macaque visual cortex and macaque frontal regions 44 and 45 (Chen et al., 2022). To test the capacity of our resource to identify novel putative areal border genes, we calculated average gradients of all genes across the boundary between mesial temporal parahippocampal gyrus (Perirhinal Ectorhinal cortex, PeEc) and the fusiform gyrus (area TF) for which there is openly available ISH data (https://human.brain-map.org/ish/search). Limiting analyses to those genes for which ISH was available, the two genes exhibiting the largest gradient in either direction (four in total) were selected. The ISH was visually inspected for the presence area-like features in gene expression. For quantitative support, the cortex in each ISH image was manually segmented over the area of interest. The pixel-wise transverse distance along the cortical segmentation from left to right was calculated and subdivided into 200 equally spaced columns, spanning from pial to white matter surface. Staining intensity was averaged across each column. For each column, we computed the t-statistic between columns to the right and left, and identified the column with the largest t-statistic as the location of the putative interareal boundary.

We used the spun+interpolated null to test whether peaks in gene gradient could be driven primarily by local folding morphology impacting non-uniform interpolation. We quantified the average gradient for all genes along the V1-V2 border in the atlas, as well as for 10 iterations of the atlas where the samples were spun prior to interpolation. We computed the median gradient magnitude for the 20 top-ranked genes for each (Fig S2d).

We benchmarked DEMs against regional differences in cellular measures of cortical organization from single nucleus RNA-sequencing studies (snRNA-seq). Specifically, we correlated regional differences in the estimated proportion of 16 neuronal subtypes across 6 cortical regions(Lake et al., 2016) with regional DEM estimates for the mean expression of provided markers for these cell types(Lake et al., 2016). The test statistic was tested against a null distribution generated through spinning and resampling the cell marker DEM estimates (***Table 1***). Given the observed correspondence between regional cellular proportions and regional expression of cell marker sets, we used more recently-generated reference cell-markers from the Allen Institute for Brain Sciences(Bakken et al., 2021; Hodge et al., 2019; Tasic et al., 2016) to generate DEMs for 11 of 14 major cell subclasses in the mammalian cortex (6 neuronal types shown in **Fig 1h**, all 11 used for TD peak enrichment analysis **Fig S3h**). Three markers were excluded due to absence in the original dataset or low gene-predictability (r<0.2, **Fig S1c**).

We benchmarked DEMs against orthogonal spatially dense measures of cortical through the following comparisons: (i) Layer IV thickness values from the 3D BigBrain atlas of cortical layers(Wagstyl et al., 2020) vs. the average DEM for later IV marker genes(He et al., 2017; Maynard et al., 2021) (**Table S2**); (ii) motor-associated areas of the cortex from multimodal in vivo MRI(Glasser et al., 2016), vs. the average DEM for two marker genes (ASGR2, CSN1S1) of Betz cells, which are giant pyramidal neurons that output from layer V of the human motor cortex(Bakken et al., 2021); (iii) an in vivo neuroimaging map of the T1/T2 ratio measuring of intracortical myelination(Glasser and Van Essen, 2011) vs. the DEM for Myelin Basic Protein; and, (iv) regional cortical thinning from in vivo sMRI data in Alzheimer disease patients with the APOE E4 (OASIS-3 dataset(LaMontagne et al., 2019), see ***MRI Data Processing*** below) vs. the APOE4 DEM. For all four of these comparisons, alignment between maps was quantified and test for statistical significance using a strict spin-based spatial permutation method that controls for spatial autocorrelation in cortical data ((Alexander-Bloch et al., 2018)methods on statistical testing of pairwise cortical maps can be found in ***Table 1***).

## 3. Characterizing the topography of DEMs

### a. Transcriptomic distinctiveness (TD) and principal component analysis (Fig 2a-c)

Transcriptomic distinctiveness (TD) of each cortical vertex was calculated as the mean of the absolute DEM value for all genes (**Fig 2a**). Statistically significant peaks in TD, driven by convergence of extreme values across multiple genes, were identified as follows. The DEM for each gene was independently spun and TD was recalculated at each vertex over 1000 sets of gene-level DEM permutations (Alexander-Bloch et al., 2018). The maximum vertex TD value for each permuted TD map was recorded and the 95th percentile value across the 1000 permutations was taken as a threshold value. This threshold represents the maximum TD one would expect in the absence of concentrated colocalisations of extreme expression signatures, and areas above this threshold were annotated as TD peaks. To disambiguate TD peaks that are spatially coalescent but potentially driven by extreme values of heterogeneous gene sets within different regions, we concatenated all suprathreshold TD vertices into a single vertex*gene matrix and vertices in this matrix were clustered based on their expression signatures.

Intervertex correlation of gene rankings were calculated and the matrix was clustered using a gaussian mixture model. Bayesian information criterion was used to identify the optimum number of clusters (k=6) from a range of 2-18. Automated labels to localize TD peaks were generated based on their intersection with a reference multimodal neuroimaging parcellation of the human cortex(Glasser et al., 2016). Each TD was given the label of the multimodal parcel that showed greatest overlap (**Fig 2b**).

The TD map was assessed for reproducibility through three approaches. First the 6-subject cohort was subdivided into pairs of triplets, for which there are 10 unique combinations. For each combination, independent TD maps were computed for each triplet and compared between triplets (**Fig S3a**). Second, for the full 6-subject cohort genes were grouped into deciles according to the reproducibility of their spatial patterns in independent sub-cohorts (**Fig S1c**). For each decile of genes a TD map was computed and compared to the TD map from the remaining 90% of genes (**Fig S3b**). Third, to assess whether the covariance in spatial patterning across genes could be a result of mesh-associated structure introduced through interpolation and smoothing, TD maps were recomputed for the spun+interpolated null datasets and compared to the original TD map (**Fig S3c)**.

The cortical regions defined by TD peaks were annotated according to their spatial overlap with the 24 cortical cell marker expression DEMs used in **Fig 1g,h** (Bakken et al., 2021; Hodge et al., 2019; Lake et al., 2016). To establish that cell maps were aligned with TD peaks, we first tested whether the vertex with the highest DEM value for each cell map overlapped with a TD peak and compared the number of overlapping cells to a null distribution created through spinning the TD peaks independently 1000 times. We then identified the cell types whose expression most closely aligned with each TD peak, comparing mean TD expression with a null distribution generated through spinning the peaks 1000 times (**Fig S3h**). TD peaks were also annotated for their functional activations using the meta-analytic Neurosynth database (see **Map annotations** below).

Gene sets characterizing TD peaks were identified as follows. At the vertex with the highest TD value within a peak region, the 95th centile TD value across genes was selected as a threshold. Genes with z-scored expression values above this threshold or below its inverse were selected, allowing TD peaks to have asymmetric length gene lists for high and low-expressed genes (**Table S3**). These TD gene lists were submitted to a Gene Ontology (GO) enrichment analysis pipeline (see ***Gene-set based annotations*** below).

To contextualize the newly-described TD peaks using previously-reported principal components (PCs) of human cortical gene expression, we computed the first 5 PC of gene expression in our full DEM library. The percentage of variance explained by each PC was calculated and compared to a null threshold derived through fitting PCs to a permuted null given by 1000 random spatial rotations of gene-level DEMs (**Fig S3d**). Taking the gene-level loadings from the first 3 PCs (**Fig S3e**), each vertex could be positioned in a 3D PC space based on its expression signature and also be colored based on its membership of a TD peak - thereby visualizing the position of TD peaks relative to the dominant spatial gradients of transcriptomic variation across the cortex (**Fig 2c**).

The assignment of TD regions as “peaks” implies a rapid emergence of the TD signature surrounding the peak boundaries, which we formally assessed by cortex-wide analysis of local tangential changes in gene expression (see “***Local Gradient Analysis***” below), and a spatially fine-grained comparisons of the physical vs. transcriptional distance between cortical regions. In the latter of these two analytic approaches, a rapid “border-like” onset of TD features would appear as (i) TD regions showing a greater transcriptional distance from other cortical regions than would be expected from their physical distance from other cortical regions, and (ii) this disparity emerging sharply surrounding the peak. To achieve this test, we first quantified the geodesic physical distance and Euclidean transcriptomic distance between pairs of vertices. For computational tractability, we limited this analysis to a subsample of vertices, choosing central vertices from ROIs in a parcellation with 500 approximately evenly sized parcels(Schaefer et al., 2018). We fit a linear generalized additive model to the data - predicting transcriptomic distance from geodesic distance - and calculated the residuals for each inter-vertex edge (**Fig S3f**). For each sampled vertex we averaged these residuals and mapped them back to the surface to visualize cortical areas that were transcriptomically more distinctive than their physical distance to other areas would predict (**Fig S3g**).

### b. Relating adult TD peaks to fetal gene expression (Fig S3k)

We sought to establish whether the regional expression signatures characterizing TD peaks were present early in fetal development. This goal required measures of gene expression from multiple regions across the fetal cortical sheet, which are provided by the Allen Institute from Brain Sciences fetal laser micro-dissection microarray dataset(Miller et al., 2014). In each samples’ fetal brain, this dataset represents approximately 25 cortical brain regions tangentially, and radially 7 transient fetal layers/compartments radially: Subpial granular zone (SG), marginal zone (MZ), outer and inner cortical plate (grouped together as CP), subplate zone (SP), intermediate zone (IZ), outer and inner subventricular zone (grouped together as SZ), and ventricular zone (VZ).

Probe-level data measures of gene expression for the two PCW21 donors in the AHBA fetal LMD microarray dataset were downloaded from (https://www.brainspan.org/static/download.html) - providing log2-transformed measures of gene expression for 58,692 probes in each of 536 tissue samples across both donors (**Table S1**). Preprocessing and normalization of these probe level gene expression values was implemented as detailed by the Allen Institute for Brain Science White Paper (https://help.brain-map.org/download/attachments/3506181/Prenatal_LMD_Microarray.pdf). Probe-level expression values were averaged for each gene to yield a single gene*sample expression matrix for each donor, which was filtered to include only cortical samples. Gene expression values were scaled across samples within each donor, and scaled gene expression values were compiled for the set of 235 cortical regions that was common to both donor datasets. We averaged scaled regional gene expression values between donors per gene, and filtered for genes in the fetal LDM dataset that were also represented in the adult DEM dataset - yielding a single final 20,476*235 gene-by-sample matrix of expression values for the human cortex at 21 PCW. Each TD peak region was then paired with the closest matching cortical label within the fetal regions. This matrix was then used to test if each TD expression signature discovered in the adult DEM dataset (**Fig 2**, **Table 3**) was already present in similar cortical regions at 21 PCW.

The analysis of fetal regional patterning of TD peak gene sets was carried out as follows (**Fig S3k**). For a given TD peak, the significantly enriched genes for that peak (see above for definition of these gene sets) were identified in the fetal dataset and averaged at each fetal sample - capturing how highly expressed the TD signature was in each fetal sample. Next, we identified all samples in the fetal expression dataset that originated from regions underlying the TD peak, and defined these as the “fetal target region set” for that TD region (i.e. occipital samples in the fetal brain were the fetal target region set for analysis of gene enriched in the adult occipital TD region). We ranked all fetal samples by their mean expression of the TD marker set, and normalized these ranks to between 0 (TD markers most highly expressed) and 1 (TD markers most lowly expressed). Normalization was done to adjust for varying numbers of areas recorded per compartment. This ranking enabled us to compute the median rank of the fetal target region set, and test if this was significantly lower compared to a null distribution of ranks from random reassignment of the fetal target region set labels across all fetal samples. Within this analytic framework, a statistically significant test means that the adult TD signature is significantly localized to homologous cortical regions at 21 PCW fetal life (**Fig S3k**). We repeated this procedure for each adult TD.

### c. Local gradient analysis (Fig 2e-g)

Spatially dense expression maps enabled the calculation of a vector describing the first spatial derivative - i.e. the local gradient - of each gene’s expression at each vertex. These vectors describe both the orientation and the magnitude of gene expression change.

Averaging these gene-level magnitude estimates across genes provided a vertex-level summary map of the magnitude of local expression changes in our full DEM library (**Fig 2e**). Regions with a significantly high average expression gradient were identified using a similar spatial permutation procedure as described for the identification of TD peaks. Briefly, the DEM gradient map for each gene was independently spun and an average expression gradient magnitude was recalculated at each vertex over 1000 sets of these spatial permutations(Alexander-Bloch et al., 2018). For each permutation we recorded the maximum vertex-level average expression gradient value, and the 95th percentile value of these maximums across the 1000 permutations was taken as a threshold value. Vertices with observed average expression gradient values above this threshold represented cortical regions of significantly rapid transcriptional change (**Fig S3j**).

The principal orientation of gene expression change at each vertex was calculated considering the vectors describing gene expression gradients - thereby providing a single summary of local gene expression gradients that considers both direction and magnitude. Principal component analysis (PCA) of gene gradient vectors was used to calculate the primary orientation of gene expression change at each vertex (**Fig 2e**) and the percentage of orientation variance accounted for by this principal component (**Fig 2e, Fig S3i**). Gene-level PC weights for each vertex were stored for subsequent analyses, including alignment with folds and functional ROIs (**Fig 2f & g**, see annotational analyses below).

The rich DEM expression gradient information described above was applied in three downstream analyses. First, we used these resources to detail the emergence of TD expression signatures within the cortical sheet - focusing on all vertices that had been identified to show a significantly elevated mean expression gradient. Specifically, we ranked genes at these vertices by their loadings onto the 1st PC of gene expression gradients at each vertex, and correlated these rankings with the rankings of genes by the expression at each TD peak vertex. This vertex-level correlation score - which quantifies how closely the gene expression gradient at a given vertex resembles that expression signature of a given TD peak - was regenerated for each of the 6 TD peaks (colors, **Fig S3j**). In each of these 6 maps, we were also able to plot the principal orientations of expression change at the vertex-level (red lines, **Fig S3i**) to ask if gradients of expression change for a given TD signature were spatially oriented towards the TD in question.

Second, we used the principal orientation of expression change at each vertex to assess whether local transcriptomic gradients were aligned with the orientation of cortical folding patterns. Orientation of cortical folds was calculated using sulcal depth and cortical curvature (Xia et al., 2018). Gradient vectors for sulcal depth describe the primary orientation of cortical folds on the walls of sulci, while gradient vectors of cortical curvature better describe the orientation at sulcal fundi and gyral crowns. These two gradient vector-fields were combined and smoothed with a 10mm FWHM gaussian kernel to propagate the vector field into plateaus e.g. at large gyral crowns where neither sulcal depth nor curvature exhibit reliable gradients. The folding orientation vectors were calculated with reference to a 2D flattened cortical representation for statistical comparison with the gradient vectors derived from gene expression maps (**Fig 2f**). At each vertex, the minimum angle was calculated between the folding orientation vector and gene expression gradient vector. Aligned vector maps exhibit positive skew, with angles tending towards zero. Therefore the skewness of the distribution of angles across all vertices was calculated, and to test for significance, folding and expression vector maps were spun relative to one another 1000 times, generating a null distribution of skewness values against which the test-statistic was compared (**Table 1**). A similar analysis was applied to test the association between module eigenmap gradient vectors and cortical folding (see ***WGCNA section*** below).

**Table 1.**
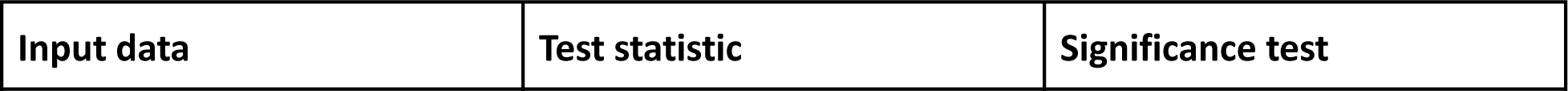
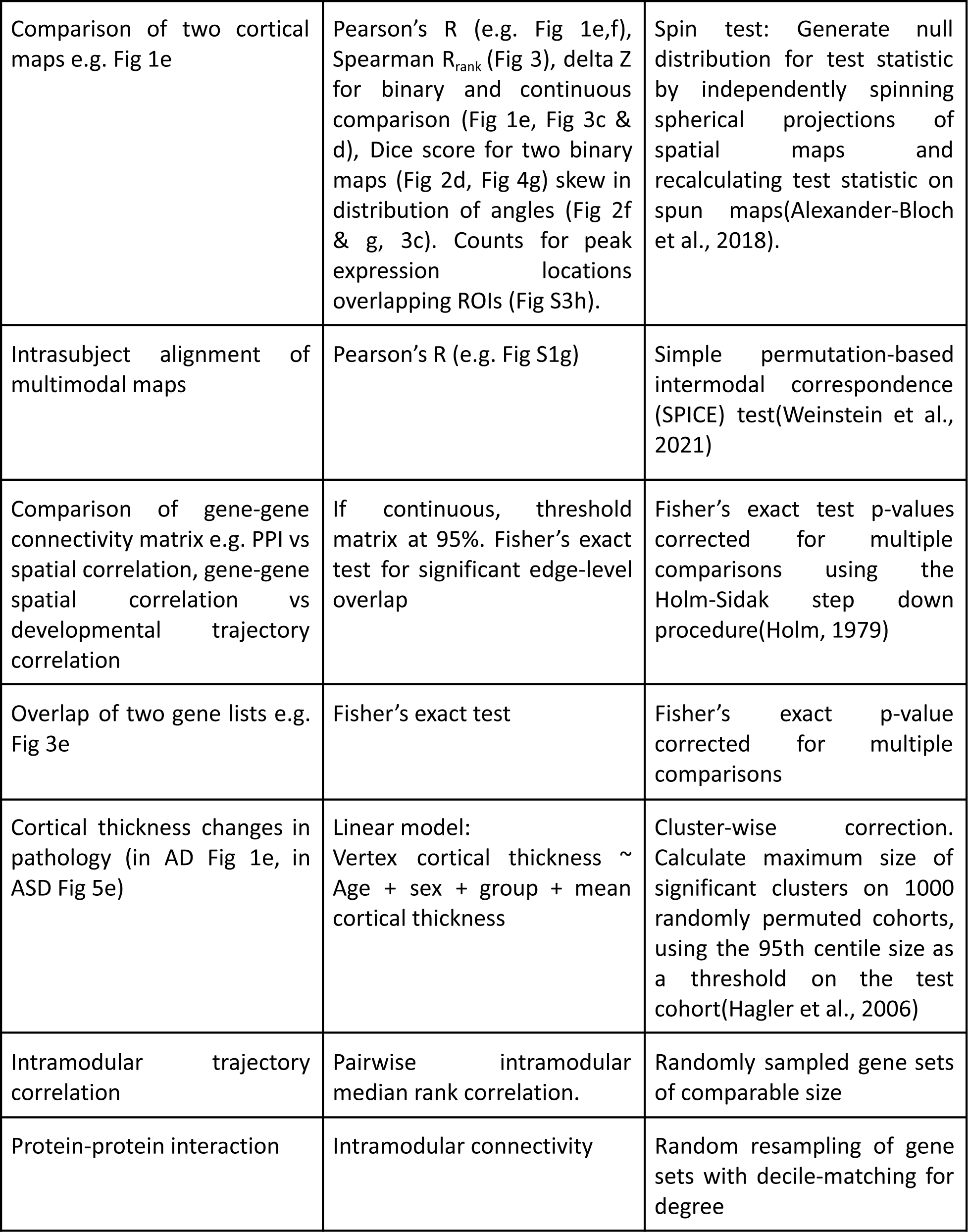
Statistical tests used to compare spatial maps and gene sets derived from the Allen Human Brain Atlas with independent multiscale neuroscientific resources.

Third we sought to quantify the alignment between cortical expression gradients and cortical areas as defined by multimodal imaging. Orientation of each MRI multimodal parcel ROI from Glasser et al(Glasser et al., 2016), was calculated taking the coordinates for all vertices within a given ROI. Principal Component Analysis of coordinates was used to identify the short and long axis of the ROI object. The vector describing the short axis was taken for comparison with mean of expression gradient vectors for vertices in the same ROI. For each ROI, the minimum angle was calculated and the skewness of the angles across all ROIs was calculated and compared to a null distribution created through spinning maps independently 1000 times, recalculating angles and their skewness (**Fig 2g**).

### d. Weighted Gene Co-expression Network Analysis (WGCNA) (Fig 3a-c)

Genes were clustered into modules for further analysis using WGCNA(Langfelder and Horvath, 2008). Briefly, gene-gene cortical spatial correlations were calculated across all vertices to generate a single square 20,781*20,781 signed co-expression matrix. This co-expression matrix underwent “soft-thresholding”, raising the values to a soft power of 6, chosen as the smallest power where the resultant network satisfied the scale-free topology model fit of r^2^>0.8(Zhang and Horvath, 2005). Next, a similarity matrix was created through calculating pairwise topological overlap, assessing the extent to which genes share neighbors in the network(Yip and Horvath, 2007). The inverse of the topological overlap matrix was then clustered using average linkage hierarchical clustering, with a minimum cluster size of 30 genes. The eigengene for each module is the first principal component of gene expression across vertices, and provides a single measure of module expression at each vertex (hence, “eigenmap”). As per past implementation of WGCNA, pairs of modules with eigengene correlations above 0.9 were merged. These procedures defined a total of 23 gene co-expression modules ranging in size from 77-3725 genes, and a single set of unconnected genes (gray module 265 genes). We filtered the gray module from further analysis, as well as all 6 other modules that were also statistically significantly enriched for NCExp genes (**Table S4**, Fisher’s test, all p<0.0001) - leaving a total of 16 modules for downstream analysis (**Table S4**). To assess the extent to which eigenmaps captured highly reproducible features of cortical organization, for each decile of genes, DEMs were correlated with their module eignmaps recomputed from the remaining 90% of genes. (**Fig S4a**).

Each WGCNA module could be visualized as a cortical eigenmap, and eigenmap gradient - on the TD terrain, or inflated cortical (**Fig 3a**). The eigenmap gradient for each module provides a vertex-level measure for the magnitude of change in module expression at each vertex, as well as a vertex-level orientation of module expression change - calculated as described in ***Local Gradient Analysis*** above. These anatomical representations of each WGCNA module are amenable to spatial comparison with any other cortical map through spatial permutations(Alexander-Bloch et al., 2018) (see ***Annotational analyses*** below). Each WGCNA module is also defined as a gene set, which is amenable to standard gene-set based enrichment analysis (see ***Annotational analyses*** below). WGCNA modules can each also be represented as a ranked list of all genes - based on gene-level kME scores for each module, which are the cross-vertex correlation between a gene’s DEM map and a module’s eigenmap.

## 4. Multiscale annotation of WGCNA modules (Fig 3c,d)

We used multiple open neuroimaging and genomic datasets to systematically sample diverse levels of cortical organization and achieve a multiscale annotation of WGCNA modules. All gene sets used in enrichment analysis are detailed in **Table S2**.

### a. Map-based annotations

#### MRI-derived maps of cortical function

Functional annotations of the cortex were carried out using two independent functional MRI (fMRI) resources - one based on resting state fMRI (rs-FMRI)(Yeo et al., 2011), and one using task-based fMRI(Rubin et al., 2017; Yarkoni et al., 2011). Resting state functional connectivity networks were taken from(Yeo et al., 2011), which divides the cortex into seven coherent functional networks through surface-based clustering of resting state fMRI into: visual, somatomotor, dorsal attention, ventral attention, frontoparietal control, limbic and default networks. We used spin-based spatial permutation testing to test for networks in which WGCNA eigenmap expression was significantly elevated (**Fig 3c, see *Table 1***).

For task fMRI-driven functional annotation of the cortex, we drew on meta-analytic maps of cortical activation from *Neurosynth(Rubin et al., 2017; Yarkoni et al., 2011)*. Briefly, over 11,000 functional neuroimaging studies were text-mined for papers containing specific terms and associated activation coordinates(Yarkoni et al., 2011). Secondary analyses generated activation maps for 30 topics spanning a range of cognitive domains (Rubin et al., 2017). Topic activation maps were intersected with cortical surface meshes and thresholded to identify vertices with an activation value above 0. Example topics included “motor, cortex, hand” and “social, reasoning, medial prefrontal cortex” (**Fig 3d**). Topics were excluded if intersected cortical maps indicated activation in fewer than 1% of cortical vertices. Topic maps were used to annotate TD peaks (**Fig 2d**) - identifying for each ROI, the 2 topics with the highest Dice overlap. Topic maps also served as an independent validation of selected WGCNA eigenmaps (**Fig 2d**, ***Table 1***). Topic maps from *Neurosynth* were also used to provide an orthogonal validation of observed resting state network enrichments from Yeo et al (**Fig 3c**) for M2 and M12: mean eigenmap expression for module M2 and M12 was calculated for *Neurosynth* topic maps and assessed for statistical significance using spin-based permutations (**Fig 3d**, ***Table 1***).

#### MRI-derived maps of cortical structure

Cortical thickness and T1/T2 “myelin” maps were taken from the Human Connectome Project average(Glasser et al., 2016). Spatial correlations were calculated across all vertices with each WGCNA module eigenmap, and assessed for statistical significance using spin-based permutations (**Fig 3c, see *Table 1***).

#### Orientation of cortical folds

We used the orientation of expression change at each vertex to assess whether local eigenmap gradients were aligned with the orientation of cortical folding patterns, mirroring the analysis described above (**Fig S4b,** see ***Local Gradient Analysis***).

#### Inter-eigenmap correlations

We tested the pairwise spatial correlation between pairs of module eigenmaps. Statistical significance was assessed using a null distribution of correlation matrices through independently spinning eigenmaps and recalculating correlations, and correcting for multiple comparisons (**Fig S4c,** see ***Table 1***).

### b. Gene-set based annotations

#### GO enrichment

Gene Ontology Enrichment Analysis (see ***Table 1*** below) were carried out on gene sets of interest, testing for enrichment of Biological Processes and Cellular Compartment, using the GOATOOLS python package(Klopfenstein et al., 2018). Where multiple gene lists were assessed simultaneously (e.g. for TD peak gene lists or WGCNA gene sets), correction for multiple comparisons was carried out by dividing the p<0.05 threshold for statistical significance by the number of tests (i.e. for 16 module p<0.05/16). To facilitate summary descriptions of multiple significant GO terms, terms were hierarchically clustered based on semantic similarity (Resnik, 1995) and representative terms were selected based on biological specificity (i.e. depth within the gene ontology tree) and magnitude of the enrichment statistic (***Fig 3d, Table S2***).

#### Layer marker gene sets and in situ hybridisation validation

We sought to assess the extent to which convergent spatial patterns of gene expression indicate convergent laminar and cellular features. Marker genes for each cortical layer were defined as the union of layer-specific marker genes from two comprehensive transcriptomic studies of layer-dependent gene expression sampling prefrontal cortical regions(He et al., 2017; Maynard et al., 2021). He et al., took human cortical samples from the prefrontal cortex, corresponding to areas BA 9, 10 & 46. Samples were sectioned into cortical depths and underwent RNAseq to identify 4131 genes exhibiting layer-dependent expression. Maynard et al., took samples from the dorsolateral prefrontal cortex and carried out spatial snRNAseq to identify 3785 genes enriched in specific cortical layers. These independent resources were combined for laminar enrichment analyses (i.e. we took each layer’s marker genes to be the union of layer genes defined in Maynard et al and He at al). WGCNA module genes were tested for laminar enrichment using Fisher’s exact test, correcting for multiple comparisons (**Fig 3c**, see ***Table 1***). Independent validation of laminar associations of candidate genes identified through the above marker lists were carried out using in situ hybridisation (ISH) data from the Allen Institute(Zeng et al., 2012). For selected modules, we identified the highest kME genes represented within the ISH dataset. For each of these genes, the highest quality sections were downloaded, and the cortical ribbon was manually segmented. Equivolumetric estimates of cortical depth were generated and profiles of depth-dependent staining intensity were generated(Huber et al., 2021). Accompanying approximate cytoarchitectonic layer thickness estimations were derived from BigBrain and used to describe the laminar location of ISH peaks(Wagstyl et al., 2020) (***Fig 3d***).

#### Adult cortical cell type marker gene sets

Cell marker gene sets were compiled from multiple snRNAseq datasets, sampling a wide variety of cortical areas covering occipital, temporal, frontal, cingulate and parietal lobes(Darmanis et al., 2015; Habib et al., 2017; Hodge et al., 2019; Lake et al., 2018, 2016; Li et al., 2018; Ruzicka et al., 2021; Velmeshev et al., 2019; Zhang et al., 2016). To integrate across differing subcategories, cell subtype marker lists were grouped into the following cell classes according to their designated names: excitatory neurons, inhibitory neurons, oligodendrocytes, astrocyte, oligodendrocyte precursor cells, microglia and endothelial cells. Marker lists for each of these cell classes represented the union of all subtypes assigned to the category. Cells not fitting into these categorisations were excluded. WGCNA module genes were tested for cell class marker enrichment using Fisher’s exact test, correcting for multiple comparisons (**Fig 3c**, see ***Table 1***).

#### Fetal cortical cell type marker gene sets

Fetal cell marker gene lists were taken from Polioudakis et al(Polioudakis et al., 2019). WGCNA module genes were tested for cell class marker enrichment using Fisher’s exact test, correcting for multiple comparisons (**Fig 3c**, see ***Table 1***).

#### Compartments and SynGO

Cellular compartment gene lists were taken from the COMPARTMENTS database(Binder et al., 2014), which identifies subcellular localisation of marker genes based on integrated information from the Human Protein Atlas, literature mining and GO annotations. Examples of cellular compartments include nucleus, plasma membrane and cytosol. An additional compartment list for neuronal synapse was generated by collapsing all genes in the manually curated SynGO dataset(Koopmans et al., 2019). WGCNA module genes were tested for cell compartment gene set enrichment using Fisher’s exact test, correcting for multiple comparisons (**Fig 3c**, see **Table 1**).

#### PPI network

Protein-protein interactions were derived from the STRING database(Szklarczyk et al., 2019). Physical direct and indirect protein-protein interactions were considered. We tested for enrichment of protein-protein interactions for proteins coded by genes within WGCNA modules. The median number of intramodular connections was compared to a null distribution of median modular connectivity derived from 10000 randomly resampled modules with the same number of genes. Gene resampling was restricted within deciles defined by the degree of protein-protein connectivity.

#### Developmental peak epoch

Peak developmental epochs for genes were extracted from(Werling et al., 2020). Briefly, bulk transcriptomic expression values were measured from DLPFC samples across development (6 PCW to 20 years), fitting developmental trajectories to each gene. Genes were categorized according to developmental epoch in which their expression peaked. For descriptive purposes, epochs were renamed as 1: “early fetal” [“fetal”, 8 postconception weeks (PCW) - 24 PCW], 2: late fetal transition (“perinatal”, 24 PCW - 6 months postnatal) and 3: “postnatal” (>6 months). Genes associated with WGCNA modules were tested for enrichment correcting for multiple comparisons across 16 modules.

#### Developmental trajectories

Gene-specific developmental trajectories were generated for the cortical samples from(Li et al., 2018). Briefly, in this study bulk transcriptomic expression values were measured from brain tissue samples taken from individuals aged between 5 PCW and 64 years old. In our analysis, samples were filtered for cortical ROIs and restricted to post 10 PCW due to lack of samples before this time-point. Ages were log transformed and Generalized Additive Models were fit to each gene to generate an estimated developmental trajectory. To compute trajectory correlations between genes, we first resampled expression trajectories at 20 equally spaced time points (in log time), and then z-normalized these values per gene (using the mean and standard deviation of each trajectory). We then calculated expression trajectory Pearson correlations between each pair of genes in this dataset, and used these to determine if the spatially co-expressed genes defining each WGCNA module also showed significant temporal co-expression. To achieve this test, we calculated the median temporal co-expression (correlation in expression trajectories) for each WGCNA module gene set, and compared this to null median co-expression values for 1000 randomly resampled gene sets matching module size. Mean trajectories of genes in each module were calculated to visualize the developmental expression pattern of each module (**Fig S4d**).

#### Fetal compartmental analysis

We used the 21 PCW fetal microarray data processed for analysis of TD peaks (see ***Relating adult TD peaks to fetal gene expression*** above, **Fig S3k**)(Miller et al., 2014), to generated marker gene sets for each of 7 transient fetal cortical compartments: subpial granular zone (SG), marginal zone (MZ), outer and inner cortical plate (grouped together as CP), subplate zone (SP), intermediate zone (IZ), outer and inner subventricular zone (grouped together as SZ), and ventricular zone (VZ). We collapsed 21 PCW cortical expression data into compartments by averaging expression values across cortical regions for each compartment because compartment differences are known to explain the bulk of variation in cortical expression within this dataset (24%(Miller et al., 2014)). The top 5% expressed genes for each of the 7 fetal compartments was taken as the compartment marker set and used for enrichment analysis of WGCNA modules with Fisher’s exact test, correcting for multiple comparisons (see ***Table 1***, **Fig 3c**).

#### Reproducibility of genes driving enrichment analyses

We calculated gene-level spatial reproducibilities for the union of all genes contributing to significant neurobiological enrichments of WGCNA modules. This was compared to a null distribution, randomly resampling the same number of genes from all those considered in the enrichment analyses.

## 5. Combining gene-set based annotations of the cortical sheet (Fig 3e, Fig S3d)

Our observation that many WGCNA modules showed statistically-significant enrichment for diverse gene sets that could span different spatial scales (e.g. layers and organelles) or temporal epochs (e.g. fetal and adult cortical features) (**Fig 3c**) suggested a potential sharing of marker gene across these diverse sets. To test this idea, and characterize potential biological themes reflected by these shared marker genes, we carried out pairwise enrichment analyses between all annotational gene lists (**Fig 3e**). Gene lists used for enrichment analysis of WGCNA modules for cortical layers, adult cells, cellular compartments, fetal cells, developmental peak epochs and fetal compartments, were taken for further analysis. A genelist-genelist pairwise enrichment matrix was generated. p-values above 0.1 were set to 1, to limit their contribution and p-values were converted to -log10(p). To remove isolated gene lists, all lists were ranked by their degree (edges defined as p<0.05) and the bottom 10% were excluded from further analysis. The matrix, excluding WGCNA modules, underwent Louvain clustering(Blondel et al., 2008), grouping together gene lists with similar properties. Clusters were assigned descriptive names according to their salient common features (e.g. Non-neuronal, Mature neuron, Mitotic, Myelin, Fetal GE) (**Fig S4e**). For visualization, the full matrix underwent UMAP embedding(McInnes et al., 2018), a non-linear dimensionality reduction technique assigning 2D coordinates to each gene list (**Fig 3e**), coloring gene lists by their assigned cluster along with the top 20% of edges.

## 6. Disease enrichment and ASD-based analysis of WGCNA modules

The proposed analyses above link regionally patterned cortical gene expression with macroscale imaging maps of structure and function, and microscale gene sets exhibiting laminar, cellular, subcellular and developmental transcriptomic specificity. We sought to assess whether WGCNA module gene lists capturing shared spatial and temporal features were also enriched for genes implicated in atypical brain development. We included genes identified in exome sequencing studies in neurodevelopmental disorders: autism spectrum disorder(Ruzzo et al., 2019; Satterstrom et al., 2020) (ASD), schizophrenia(Singh et al., 2020) (SCZ), severe developmental disorders(Deciphering Developmental Disorders Study, 2017) (Deciphering Developmental Disorders study, DDD) and epilepsy(Heyne et al., 2018). WGCNA module gene sets were tested for enrichment of these genes using Fisher’s test and corrected for multiple comparisons (***Table 1***, **Fig 4a**). Two modules - M12 and M15 - showed enrichment for multiple disease sets, with the ASD gene set being unique for showing enrichment in both modules. We therefore focused downstream analysis on further characterizing the enrichment of ASD genes in M12 and M15, and testing if these enrichments could predict regional cortical changes in ASD.

### a. Characterizing ASD gene enrichments in M12 and M15

#### kME analysis

To better characterize the spatially distinctive properties of genes within M12 and M15, we defined the union of genes in both modules and collated the WGCNA-defined kME scores for each gene to both M12 and M15. This provided a basis for plotting all genes by their relative membership to both modules to: quantify the proximity of each gene to each module; assess the discreteness of gene assignment to modules; and - for any provide a common space within which to project gene functions and associations with ASD (**Fig 4c**)

#### Enrichment of ASD-linked GO terms

Genes linked to two specific GO terms, “Neuronal communication” and “Gene expression regulation”, enriched amongst risk genes for Autism Spectrum Disorder in(Satterstrom et al., 2020), were separately tested for enrichment within M12 and M15 (**Fig 4d**), using a Fisher’s exact test.

#### Developmental trajectories of disease-linked modules

To characterize the distinctive temporal trajectories of M12 & M15 (see **Fig 3c**), we took gene-level trajectories (see ***Developmental trajectories*** above) and calculated the mean gene-expression trajectory of genes in each module (**Fig 4e**).

#### Independent characterisation of ASD risk genes

To assess the extent to which modules M12 & M15 captured the underlying axes of spatial patterning across all 135 ASD risk genes, we took DEMs for all 135 risk genes and independently clustered them. Pairwise co-expression was calculated for all risk gene DEMs and the resultant matrix was clustered using Gaussian mixture modeling into two clusters, C1 and C2 (**Fig S5a**). kME values were calculated for each risk gene with all WGCNA modules and averaged within each cluster. For each cluster, we then identified the WGCNA module with the highest mean kME (**Fig S5b**)

### b. Comparing M12 and M15 expression to regional changes of cortical gene expression in ASD (Fig 4f)

We mapped regional transcriptomic disruption in ASD measured from multiple cortical regions using RNA-seq data(Haney et al., 2020). This study compared bulk transcriptomic expression in ASD and control samples across 11 cortical areas, quantifying the extent of transcriptomic disruption by identifying the number of significantly differentially expressed genes in each region. Cortical areas sampled in this study were mapped to their closest corresponding area in a multimodal MRI parcellation(Glasser et al., 2016). The mean expression of M12 & M15 eigenmaps was quantified in the same cortical areas (**Fig 4f**). The test statistic, correlating eigenmap expression with the number of differentially expressed genes, was tested against a null distribution generated through spinning and resampling the eigenmaps (see **Table 1**).

### c. Comparing M12 and M15 expression to regional changes of cortical thickness in ASD (Fig 4g, h, Fig S5c)

To assess the extent to which WGCNA module eigenmaps pattern macroscale *in vivo* anatomical differences in ASD, we took the map of relative cortical thickness change in autism (see ***Preprocessing and analysis of structural MRI data*** below) and compared this to eigenmap expression patterns. M12 and M15 eigenmaps were thresholded, identifying the 5% of vertices with the highest expression. Areas of high significant thickness change were tested for overlap with areas of significant cortical thickness change using the Dice overlap compared to a null distribution of Dice scores generated through spinning the thresholded eigenmaps (see **Table 1**)

## 7. Preprocessing and analysis of structural MRI data

### a. AHBA donors

Pial and white matter cortical T1 MRI scans of the 6 AHBA donor brains were reconstructed using Freesurfer (v5.3)(Romero-Garcia et al., 2018)(see **Table S1**). Briefly, scans undergo tissue segmentation, cortical white and pial surface extraction. A mid-thickness surface, between pial and white surfaces was also created. The locations of tissue samples taken for bulk transcriptomic profiling, provided in the coordinates of the subject’s MRI were mapped to the mid-thickness surface as outlined above (see **Creating spatially dense maps of human cortical gene expression from the AHBA**). Individual subject cortical surfaces were co-registered to the fs_LR32k template surface brain using MSMSulc(Robinson et al., 2018) as part of the ciftify pipeline(Dickie et al., 2019), which warps subject meshes by non-linear alignment their folding patterns to the MRI-derived template surface. A donor-specific template surface was created through averaging the coordinates of the aligned meshes and used for analysis of cortical folding patterns used in **Alignment with reference measures of cortical organization.** Pial, Inflated and flattened representations of the fs_LR32k surface were used for the visualization of cortical maps throughout.

### b. OASIS (Fig 1e)

To estimate relative cortical thickness change in AD patients with the APOE E4 variant, we utilized the openly available OASIS database(LaMontagne et al., 2019). T1w MRI data collected using a Siemens Tim Trio 3T scanner and underwent cortical surface reconstruction using Freesurfer v5.3 as above. Reconstructions underwent manual quality control and correction, with poor quality data being removed. Output cortical thickness maps, smoothed at 20mm fwhm and aligned to the fsaverage template surface were downloaded via https://www.oasis-brains.org/, along with age, sex, APOE genotype and cognitive status. Subjects were included in the analysis if they had been diagnosed with AD and had at least one APOE E4 allele (n=119), or were a healthy control (n=633) (see **Table S1**). Per-vertex coefficients for disease-associated cortical thinning and significance were calculated, adjusting for age, sex and mean cortical thickness. We controlled for mean CT to identify local anatomical changes given our finding of generalized cortical thickening in AD as compared to controls in OASIS. The map of cortical thickness coefficients was then registered from fsaverage to fs_LR32k for comparison with the DEM of APOE (**Fig 1e**)(Robinson et al., 2018).

### c. ABIDE

To estimate relative cortical thickness change in ASD, MRI cortical thickness maps, generated through Freesurfer processing of 3T T1 structural MRI scans were downloaded from ABIDE, along with age, sex, site information(Di Martino et al., 2017, 2013)(**Table S1**). Multiple sites and scanners were used to acquire these data, which is known to introduce systematic biases in morphological measurements like cortical thickness. To mitigate this, we used neuroCombat which estimates and removes unwanted scanner-effects while retaining biological effects on variables such as age, sex and diagnosis(Fortin et al., 2018). Subjects with poor quality freesurfer segmentations were excluded using a threshold Euler count of 100 (ref). Cortical thickness change in ASD relative to controls was calculated adjusting for age, sex and mean cortical thickness. Neighbor-connected vertices exhibiting significant cortical thickness change (p<0.05) were grouped into clusters. A null distribution of cluster sizes was generated using 1000 random permutations of the cohort, storing the maximum significant cluster size for each permutation. The 95th percentile cluster size was used as a threshold for removing test clusters that could have arisen by chance(Hagler et al., 2006). Output coefficient and cluster maps were registered from fsaverage to fs_LR32k and compared with the M12 and M15 eigenmaps as described above.

## Acknowledgements

The authors would like to thank all the participants and their families for their generous involvement in this study.

## Data availability

The cortical dense expression and gradient maps of 20,781 genes and ∼30k vertices that support the findings of this study are available at https://figshare.com/s/82c8f6ebda38af670cd1. Scripts to download, visualize and analyze MAGICC are available at https://github.com/kwagstyl/MAGICC.

## References

Alexander-Bloch AF, Shou H, Liu S, Satterthwaite TD, Glahn DC, Shinohara RT, Vandekar SN, Raznahan A. 2018. On testing for spatial correspondence between maps of human brain structure and function. Neuroimage 178:540–551.

Arnatkeviciute A, Fulcher BD, Fornito A. 2019. A practical guide to linking brain-wide gene expression and neuroimaging data. Neuroimage 189:353–367.

Bakken TE, Jorstad NL, Hu Q, Lake BB, Tian W, Kalmbach BE, Crow M, Hodge RD, Krienen FM, Sorensen SA, Eggermont J, Yao Z, Aevermann BD, Aldridge AI, Bartlett A, Bertagnolli D, Casper T, Castanon RG, Crichton K, Daigle TL, Dalley R, Dee N, Dembrow N, Diep D, Ding S-L, Dong W, Fang R, Fischer S, Goldman M, Goldy J, Graybuck LT, Herb BR, Hou X, Kancherla J, Kroll M, Lathia K, van Lew B, Li YE, Liu CS, Liu H, Lucero JD, Mahurkar A, McMillen D, Miller JA, Moussa M, Nery JR, Nicovich PR, Niu S-Y, Orvis J, Osteen JK, Owen S, Palmer CR, Pham T, Plongthongkum N, Poirion O, Reed NM, Rimorin C, Rivkin A, Romanow WJ, Sedeño-Cortés AE, Siletti K, Somasundaram S, Sulc J, Tieu M, Torkelson A, Tung H, Wang X, Xie F, Yanny AM, Zhang R, Ament SA, Behrens MM, Bravo HC, Chun J, Dobin A, Gillis J, Hertzano R, Hof PR, Höllt T, Horwitz GD, Keene CD, Kharchenko PV, Ko AL, Lelieveldt BP, Luo C, Mukamel EA, Pinto-Duarte A, Preissl S, Regev A, Ren B, Scheuermann RH, Smith K, Spain WJ, White OR, Koch C, Hawrylycz M, Tasic B, Macosko EZ, McCarroll SA, Ting JT, Zeng H, Zhang K, Feng G, Ecker JR, Linnarsson S, Lein ES. 2021. Comparative cellular analysis of motor cortex in human, marmoset and mouse. Nature 598:111–119.

Binder JX, Pletscher-Frankild S, Tsafou K, Stolte C, O’Donoghue SI, Schneider R, Jensen LJ. 2014. COMPARTMENTS: unification and visualization of protein subcellular localization evidence. Database 2014:bau012.

Blondel VD, Guillaume J-L, Lambiotte R, Lefebvre E. 2008. Fast unfolding of communities in large networks. arXiv [physics.soc-ph].

Brodmann K. 1909. Vergleichende Lokalisationslehre der Grosshirnrinde in ihren Prinzipien dargestellt auf Grund des Zellenbaues. Barth.

Burt JB, Demirtaş M, Eckner WJ, Navejar NM, Ji JL, Martin WJ, Bernacchia A, Anticevic A, Murray JD. 2018. Hierarchy of transcriptomic specialization across human cortex captured by structural neuroimaging topography. Nat Neurosci 21:1251–1259.

Burt JB, Helmer M, Shinn M, Anticevic A, Murray JD. 2020. Generative modeling of brain maps with spatial autocorrelation. Neuroimage 220:117038.

Chen A, Sun Y, Lei Y, Li C, Liao S, Liang Z, Lin F, Yuan N, Li M, Wang K, Yang M, Zhang S, Zhuang Z, Meng J, Song Q, Zhang Y, Xu Y, Cui L, Han L, Yang H, Sun X, Fei T, Chen B, Li W, Huangfu B, Ma K, Li Z, Lin Y, Liu Z, Wang H, Zhong Y, Zhang H, Yu Q, Wang Y, Zhu Z, Liu X, Peng J, Liu C, Chen W, An Y, Xia S, Lu Y, Wang M, Song X, Liu S, Wang Z, Gong C, Huang X, Yuan Y, Zhao Y, Luo Z, Tan X, Liu J, Zheng M, Li S, Huang Y, Hong Y, Huang Z, Li M, Zhang R, Jin M, Li Y, Zhang H, Sun S, Bai Y, Cheng M, Hu G, Liu S, Wang B, Xiang B, Li S, Li H, Chen M, Wang S, Zhang Q, Liu W, Liu X, Zhao Q, Lisby M, Wang J, Fang J, Lu Z, Lin Y, Xie Q, He J, Xu H, Huang W, Wei W, Yang H, Sun Y, Poo M, Wang J, Li Y, Shen Z, Liu L, Liu Z, Xu X, Li C. 2022. Global Spatial Transcriptome of Macaque Brain at Single-Cell Resolution. bioRxiv. doi:10.1101/2022.03.23.485448

Chi JG, Dooling EC, Gilles FH. 1977. Gyral development of the human brain. Ann Neurol 1:86–93.

Collins CE, Airey DC, Young NA, Leitch DB, Kaas JH. 2010. Neuron densities vary across and within cortical areas in primates. Proceedings of the National Academy of Sciences 107:15927–15932.

Collins CE, Turner EC, Sawyer EK, Reed JL, Young NA, Flaherty DK, Kaas JH. 2016. Cortical cell and neuron density estimates in one chimpanzee hemisphere. Proc Natl Acad Sci U S A 113:740–745.

Darmanis S, Sloan SA, Zhang Y, Enge M, Caneda C, Shuer LM, Hayden Gephart MG, Barres BA, Quake SR. 2015. A survey of human brain transcriptome diversity at the single cell level. Proc Natl Acad Sci U S A 112:7285–7290.

Deciphering Developmental Disorders Study. 2017. Prevalence and architecture of de novo mutations in developmental disorders. Nature 542:433–438.

de Kovel CGF, Lisgo SN, Fisher SE, Francks C. 2018. Subtle left-right asymmetry of gene expression profiles in embryonic and foetal human brains. Sci Rep 8:12606.

Dickie EW, Anticevic A, Smith DE, Coalson TS, Manogaran M, Calarco N, Viviano JD, Glasser MF, Van Essen DC, Voineskos AN. 2019. Ciftify: A framework for surface-based analysis of legacy MR acquisitions. Neuroimage 197:818–826.

Di Martino A, O’Connor D, Chen B, Alaerts K, Anderson JS, Assaf M, Balsters JH, Baxter L, Beggiato A, Bernaerts S, Blanken LME, Bookheimer SY, Braden BB, Byrge L, Castellanos FX, Dapretto M, Delorme R, Fair DA, Fishman I, Fitzgerald J, Gallagher L, Keehn RJJ, Kennedy DP, Lainhart JE, Luna B, Mostofsky SH, Müller R-A, Nebel MB, Nigg JT, O’Hearn K, Solomon M, Toro R, Vaidya CJ, Wenderoth N, White T, Craddock RC, Lord C, Leventhal B, Milham MP. 2017. Enhancing studies of the connectome in autism using the autism brain imaging data exchange II. Sci Data 4:170010.

Di Martino A, Yan C-G, Li Q, Denio E, Castellanos FX, Alaerts K, Anderson JS, Assaf M, Bookheimer SY, Dapretto M, Deen B, Delmonte S, Dinstein I, Ertl-Wagner B, Fair DA, Gallagher L, Kennedy DP, Keown CL, Keysers C, Lainhart JE, Lord C, Luna B, Menon V, Minshew NJ, Monk CS, Mueller S, Müller R-A, Nebel MB, Nigg JT, O’Hearn K, Pelphrey KA, Peltier SJ, Rudie JD, Sunaert S, Thioux M, Tyszka JM, Uddin LQ, Verhoeven JS, Wenderoth N, Wiggins JL, Mostofsky SH, Milham MP. 2013. The autism brain imaging data exchange: towards a large-scale evaluation of the intrinsic brain architecture in autism. Mol Psychiatry 19:659–667.

Figueroa RL, Zeng-Treitler Q, Kandula S, Ngo LH. 2012. Predicting sample size required for classification performance. BMC Med Inform Decis Mak 12:8. Fischl B. 2012. FreeSurfer. Neuroimage 62:774–781.

Fortin J-P, Cullen N, Sheline YI, Taylor WD, Aselcioglu I, Cook PA, Adams P, Cooper C, Fava M, McGrath PJ, McInnis M, Phillips ML, Trivedi MH, Weissman MM, Shinohara RT. 2018. Harmonization of cortical thickness measurements across scanners and sites. Neuroimage 167:104–120.

Geschwind DH, Rakic P. 2013. Cortical evolution: judge the brain by its cover. Neuron 80:633–647.

Girault JB, Donovan K, Hawks Z, Talovic M, Forsen E, Elison JT, Shen MD, Swanson MR, Wolff JJ, Kim SH, Nishino T, Davis S, Snyder AZ, Botteron KN, Estes AM, Dager SR, Hazlett HC, Gerig G, McKinstry R, Pandey J, Schultz RT, St John T, Zwaigenbaum L, Todorov A, Truong Y, Styner M, Pruett JR Jr, Constantino JN, Piven J, IBIS Network. 2022. Infant Visual Brain Development and Inherited Genetic Liability in Autism. Am J Psychiatry 179:573–585.

Glasser MF, Coalson TS, Robinson EC, Hacker CD, Harwell J, Yacoub E, Ugurbil K, Andersson J, Beckmann CF, Jenkinson M, Smith SM, Van Essen DC. 2016. A multi-modal parcellation of human cerebral cortex. Nature 536:171–178.

Glasser MF, Sotiropoulos SN, Wilson JA, Coalson TS, Fischl B, Andersson JL, Xu J, Jbabdi S, Webster M, Polimeni JR, Van Essen DC, Jenkinson M, WU-Minn HCP Consortium. 2013. The minimal preprocessing pipelines for the Human Connectome Project. Neuroimage 80:105–124.

Glasser MF, Van Essen DC. 2011. Mapping human cortical areas in vivo based on myelin content as revealed by T1- and T2-weighted MRI. J Neurosci 31:11597–11616.

Gryglewski G, Seiger R, James GM, Godbersen GM, Komorowski A, Unterholzner J, Michenthaler P, Hahn A, Wadsak W, Mitterhauser M, Kasper S, Lanzenberger R. 2018. Spatial analysis and high resolution mapping of the human whole-brain transcriptome for integrative analysis in neuroimaging. Neuroimage 176:259–267.

Gutiérrez-Galve L, Lehmann M, Hobbs NZ, Clarkson MJ, Ridgway GR, Crutch S, Ourselin S, Schott JM, Fox NC, Barnes J. 2009. Patterns of cortical thickness according to APOE genotype in Alzheimer’s disease. Dement Geriatr Cogn Disord 28:476–485.

Habib N, Avraham-Davidi I, Basu A, Burks T, Shekhar K, Hofree M, Choudhury SR, Aguet F, Gelfand E, Ardlie K, Weitz DA, Rozenblatt-Rosen O, Zhang F, Regev A. 2017. Massively parallel single-nucleus RNA-seq with DroNc-seq. Nat Methods 14:955–958.

Hagler DJ Jr, Saygin AP, Sereno MI. 2006. Smoothing and cluster thresholding for cortical surface-based group analysis of fMRI data. Neuroimage 33:1093–1103.

Haney JR, Wamsley B, Chen GT, Parhami S, Emani PS, Chang N, Hoftman GD, de Alba D, Kale G, Ramaswami G, Hartl CL, Jin T, Wang D, Ou J, Wu YE, Parikshak NN, Swarup V, Grant Belgard T, Gerstein M, Pasaniuc B, Gandal MJ, Geschwind DH. 2020. Broad transcriptomic dysregulation across the cerebral cortex in ASD. bioRxiv. doi:10.1101/2020.12.17.423129

Hansen JY, Markello RD, Vogel JW, Seidlitz J, Bzdok D, Misic B. 2021. Mapping gene transcription and neurocognition across human neocortex. Nature Human Behaviour. doi:10.1038/s41562-021-01082-z

Hartl CL, Ramaswami G, Pembroke WG, Muller S, Pintacuda G, Saha A, Parsana P, Battle A, Lage K, Geschwind DH. 2021. Coexpression network architecture reveals the brain-wide and multiregional basis of disease susceptibility. Nat Neurosci 24:1313–1323.

Hawrylycz MJ, Lein ES, Guillozet-Bongaarts AL, Shen EH, Ng L, Miller JA, van de Lagemaat LN, Smith KA, Ebbert A, Riley ZL, Abajian C, Beckmann CF, Bernard A, Bertagnolli D, Boe AF, Cartagena PM, Chakravarty MM, Chapin M, Chong J, Dalley RA, Daly BD, Dang C, Datta S, Dee N, Dolbeare TA, Faber V, Feng D, Fowler DR, Goldy J, Gregor BW, Haradon Z, Haynor DR, Hohmann JG, Horvath S, Howard RE, Jeromin A, Jochim JM, Kinnunen M, Lau C, Lazarz ET, Lee C, Lemon TA, Li L, Li Y, Morris JA, Overly CC, Parker PD, Parry SE, Reding M, Royall JJ, Schulkin J, Sequeira PA, Slaughterbeck CR, Smith SC, Sodt AJ, Sunkin SM, Swanson BE, Vawter MP, Williams D, Wohnoutka P, Zielke HR, Geschwind DH, Hof PR, Smith SM, Koch C, Grant SG, Jones AR. 2012. An anatomically comprehensive atlas of the adult human brain transcriptome. Nature 489:391–399.

Hawrylycz M, Miller JA, Menon V, Feng D, Dolbeare T, Guillozet-Bongaarts AL, Jegga AG, Aronow BJ, Lee C-K, Bernard A, Glasser MF, Dierker DL, Menche J, Szafer A, Collman F, Grange P, Berman KA, Mihalas S, Yao Z, Stewart L, Barabási A-L, Schulkin J, Phillips J, Ng L, Dang C, Haynor DR, Jones A, Van Essen DC, Koch C, Lein E. 2015. Canonical genetic signatures of the adult human brain. Nat Neurosci 18:1832–1844.

Heuer K, Toro R. 2019. Role of mechanical morphogenesis in the development and evolution of the neocortex. Phys Life Rev 31:233–239.

Heyne HO, Singh T, Stamberger H, Abou Jamra R, Caglayan H, Craiu D, De Jonghe P, Guerrini R, Helbig KL, Koeleman BPC, Kosmicki JA, Linnankivi T, May P, Muhle H, Møller RS, Neubauer BA, Palotie A, Pendziwiat M, Striano P, Tang S, Wu S, EuroEPINOMICS RES Consortium, Poduri A, Weber YG, Weckhuysen S, Sisodiya SM, Daly MJ, Helbig I, Lal D, Lemke JR. 2018. De novo variants in neurodevelopmental disorders with epilepsy. Nat Genet 50:1048–1053.

He Z, Han D, Efimova O, Guijarro P, Yu Q, Oleksiak A, Jiang S, Anokhin K, Velichkovsky B, Grünewald S, Khaitovich P. 2017. Comprehensive transcriptome analysis of neocortical layers in humans, chimpanzees and macaques. Nat Neurosci 20:886–895.

Hodge RD, Bakken TE, Miller JA, Smith KA, Barkan ER, Graybuck LT, Close JL, Long B, Johansen N, Penn O, Yao Z, Eggermont J, Höllt T, Levi BP, Shehata SI, Aevermann B, Beller A, Bertagnolli D, Brouner K, Casper T, Cobbs C, Dalley R, Dee N, Ding S-L, Ellenbogen RG, Fong O, Garren E, Goldy J, Gwinn RP, Hirschstein D, Keene CD, Keshk M, Ko AL, Lathia K, Mahfouz A, Maltzer Z, McGraw M, Nguyen TN, Nyhus J, Ojemann JG, Oldre A, Parry S, Reynolds S, Rimorin C, Shapovalova NV, Somasundaram S, Szafer A, Thomsen ER, Tieu M, Quon G, Scheuermann RH, Yuste R, Sunkin SM, Lelieveldt B, Feng D, Ng L, Bernard A, Hawrylycz M, Phillips JW, Tasic B, Zeng H, Jones AR, Koch C, Lein ES. 2019. Conserved cell types with divergent features in human versus mouse cortex. Nature 573:61–68.

Holm S. 1979. A Simple Sequentially Rejective Multiple Test Procedure. Scand Stat Theory Appl 6:65–70.

Huber L (renzo), Poser BA, Bandettini PA, Arora K, Wagstyl K, Cho S, Goense J, Nothnagel N, Morgan AT, van den Hurk J, Müller AK, Reynolds RC, Glen DR, Goebel R, Gulban OF. 2021. LayNii: A software suite for layer-fMRI. Neuroimage 237:118091.

Jo HJ, Saad ZS, Gotts SJ, Martin A, Cox RW. 2012. Quantifying agreement between anatomical and functional interhemispheric correspondences in the resting brain. PLoS One 7:e48847.

Kelley KW, Nakao-Inoue H, Molofsky AV, Oldham MC. 2018. Variation among intact tissue samples reveals the core transcriptional features of human CNS cell classes. Nat Neurosci 21:1171–1184.

Klopfenstein DV, Zhang L, Pedersen BS, Ramírez F, Warwick Vesztrocy A, Naldi A, Mungall CJ, Yunes JM, Botvinnik O, Weigel M, Dampier W, Dessimoz C, Flick P, Tang H. 2018. GOATOOLS: A Python library for Gene Ontology analyses. Sci Rep 8:10872.

Koopmans F, Van Nierop P, Andres-Alonso M, Byrnes A, Cijsouw T, Coba MP, Cornelisse LN, Farrell RJ, Goldschmidt HL, Howrigan DP, Hussain NK, Imig C, de Jong APH, Jung H, Kohansalnodehi M, Kramarz B, Lipstein N, Lovering RC, MacGillavry H, Mariano V, Mi H, Ninov M, Osumi-Sutherland D, Pielot R, Smalla K-H, Tang H, Tashman K, Toonen RFG, Verpelli C, Reig-Viader R, Watanabe K, Van Weering J, Achsel T, Ashrafi G, Asi N, Brown TC, De Camilli P, Feuermann M, Foulger RE, Gaudet P, Joglekar A, Kanellopoulos A, Malenka R, Nicoll RA, Pulido C, de Juan-Sanz J, Sheng M, Südhof TC, Tilgner HU, Bagni C, Bayés À, Biederer T, Brose N, Chua JJE, Dieterich DC, Gundelfinger ED, Hoogenraad C, Huganir RL, Jahn R, Kaeser PS, Kim E, Kreutz MR, McPherson PS, Neale BM, O’Connor V, Posthuma D, Ryan TA, Sala C, Feng G, Hyman SE, Thomas PD, Smit AB, Verhage M. 2019. SynGO: An Evidence-Based, Expert-Curated Knowledge Base for the Synapse. Neuron 103:217–234.e4.

Kravitz DJ, Saleem KS, Baker CI, Ungerleider LG, Mishkin M. 2013. The ventral visual pathway: an expanded neural framework for the processing of object quality. Trends in Cognitive Sciences. doi:10.1016/j.tics.2012.10.011

Lake BB, Ai R, Kaeser GE, Salathia NS, Yung YC, Liu R, Wildberg A, Gao D, Fung H-L, Chen S, Vijayaraghavan R, Wong J, Chen A, Sheng X, Kaper F, Shen R, Ronaghi M, Fan J-B, Wang W, Chun J, Zhang K. 2016. Neuronal subtypes and diversity revealed by single-nucleus RNA sequencing of the human brain. Science 352:1586–1590.

Lake BB, Chen S, Sos BC, Fan J, Kaeser GE, Yung YC, Duong TE, Gao D, Chun J, Kharchenko PV, Zhang K. 2018. Integrative single-cell analysis of transcriptional and epigenetic states in the human adult brain. Nat Biotechnol 36:70–80.

LaMontagne PJ, Benzinger TLS, Morris JC, Keefe S, Hornbeck R, Xiong C, Grant E, Hassenstab J, Moulder K, Vlassenko AG, Raichle ME, Cruchaga C, Marcus D. 2019. OASIS-3: Longitudinal neuroimaging, clinical, and cognitive dataset for normal aging and Alzheimer disease. bioRxiv. doi:10.1101/2019.12.13.19014902

Langfelder P, Horvath S. 2008. WGCNA: an R package for weighted correlation network analysis. BMC Bioinformatics 9:559.

Larivière S, Paquola C, Park B-Y, Royer J, Wang Y, Benkarim O, de Wael RV, Valk SL, Thomopoulos SI, Kirschner M, Lewis LB, Evans AC, Sisodiya SM, McDonald CR, Thompson PM, Bernhardt BC. 2021. The ENIGMA Toolbox: multiscale neural contextualization of multisite neuroimaging datasets. Nature Methods. doi:10.1038/s41592-021-01186-4

Lefèvre J, Pepe A, Muscato J, De Guio F, Girard N, Auzias G, Germanaud D. 2018. SPANOL (SPectral ANalysis of Lobes): A Spectral Clustering Framework for Individual and Group Parcellation of Cortical Surfaces in Lobes. Front Neurosci 12:354.

Li M, Santpere G, Imamura Kawasawa Y, Evgrafov OV, Gulden FO, Pochareddy S, Sunkin SM, Li Z, Shin Y, Zhu Y, Sousa AMM, Werling DM, Kitchen RR, Kang HJ, Pletikos M, Choi J, Muchnik S, Xu X, Wang D, Lorente-Galdos B, Liu S, Giusti-Rodríguez P, Won H, de Leeuw CA, Pardiñas AF, BrainSpan Consortium, PsychENCODE Consortium, PsychENCODE Developmental Subgroup, Hu M, Jin F, Li Y, Owen MJ, O’Donovan MC, Walters JTR, Posthuma D, Reimers MA, Levitt P, Weinberger DR, Hyde TM, Kleinman JE, Geschwind DH, Hawrylycz MJ, State MW, Sanders SJ, Sullivan PF, Gerstein MB, Lein ES, Knowles JA, Sestan N. 2018. Integrative functional genomic analysis of human brain development and neuropsychiatric risks. Science 362. doi:10.1126/science.aat7615

Llinares-Benadero C, Borrell V. 2019. Deconstructing cortical folding: genetic, cellular and mechanical determinants. Nat Rev Neurosci. doi:10.1038/s41583-018-0112-2

Markello RD, Arnatkeviciute A, Poline J-B, Fulcher BD, Fornito A, Misic B. 2021. Standardizing workflows in imaging transcriptomics with the abagen toolbox. Elife 10. doi:10.7554/eLife.72129

Markello RD, Misic B. 2021. Comparing spatial null models for brain maps. Neuroimage 236:118052.

Maynard KR, Collado-Torres L, Weber LM, Uytingco C, Barry BK, Williams SR, Catallini JL, Tran MN, Besich Z, Tippani M, Chew J, Yin Y, Kleinman JE, Hyde TM, Rao N, Hicks SC, Martinowich K, Jaffe AE. 2021. Transcriptome-scale spatial gene expression in the human dorsolateral prefrontal cortex. Nat Neurosci 24:425–436.

McInnes L, Healy J, Melville J. 2018. UMAP: Uniform Manifold Approximation and Projection for Dimension Reduction. arXiv [statML].

Menassa DA, Gomez-Nicola D. 2018. Microglial Dynamics During Human Brain Development. Front Immunol 9:1014.

Mesulam MM. 1998. From sensation to cognition. Brain 121 (Pt 6):1013–1052.

Miller JA, Ding S-L, Sunkin SM, Smith KA, Ng L, Szafer A, Ebbert A, Riley ZL, Royall JJ, Aiona K, Arnold JM, Bennet C, Bertagnolli D, Brouner K, Butler S, Caldejon S, Carey A, Cuhaciyan C, Dalley RA, Dee N, Dolbeare TA, Facer BAC, Feng D, Fliss TP, Gee G, Goldy J, Gourley L, Gregor BW, Gu G, Howard RE, Jochim JM, Kuan CL, Lau C, Lee C-K, Lee F, Lemon TA, Lesnar P, McMurray B, Mastan N, Mosqueda N, Naluai-Cecchini T, Ngo N-K, Nyhus J, Oldre A, Olson E, Parente J, Parker PD, Parry SE, Stevens A, Pletikos M, Reding M, Roll K, Sandman D, Sarreal M, Shapouri S, Shapovalova NV, Shen EH, Sjoquist N, Slaughterbeck CR, Smith M, Sodt AJ, Williams D, Zöllei L, Fischl B, Gerstein MB, Geschwind DH, Glass IA, Hawrylycz MJ, Hevner RF, Huang H, Jones AR, Knowles JA, Levitt P, Phillips JW, Sestan N, Wohnoutka P, Dang C, Bernard A, Hohmann JG, Lein ES. 2014. Transcriptional landscape of the prenatal human brain. Nature 508:199–206.

Molnár Z, Clowry GJ, Šestan N, Alzu’bi A, Bakken T, Hevner RF, Hüppi PS, Kostović I, Rakic P, Anton ES, Edwards D, Garcez P, Hoerder-Suabedissen A, Kriegstein A. 2019. New insights into the development of the human cerebral cortex. J Anat 235:432–451.

Monier A, Adle-Biassette H, Delezoide A-L, Evrard P, Gressens P, Verney C. 2007. Entry and distribution of microglial cells in human embryonic and fetal cerebral cortex. J Neuropathol Exp Neurol 66:372–382.

Moresi L, Mather B. 2019. Stripy: A Python module for (constrained) triangulation in Cartesian coordinates and on a sphere. J Open Source Softw 4:1410.

Nieuwenhuys R, Broere CAJ. 2017. A map of the human neocortex showing the estimated overall myelin content of the individual architectonic areas based on the studies of Adolf Hopf. Brain Struct Funct 222:465–480.

O’Leary DD. 1989. Do cortical areas emerge from a protocortex? Trends Neurosci 12:400–406.

O’Leary DDM, Chou S-J, Sahara S. 2007. Area patterning of the mammalian cortex. Neuron 56:252–269.

Palomero-Gallagher N, Zilles K. 2019. Cortical layers: Cyto-, myelo-, receptor-and synaptic architecture in human cortical areas. Neuroimage 197:716–741.

Pang JC, Aquino KM, Oldehinkel M, Robinson PA, Fulcher BD, Breakspear M, Fornito A. 2023. Geometric constraints on human brain function. Nature 618:566–574.

Parikshak NN, Luo R, Zhang A, Won H, Lowe JK, Chandran V, Horvath S, Geschwind DH. 2013. Integrative functional genomic analyses implicate specific molecular pathways and circuits in autism. Cell 155:1008–1021.

Pfeifer RA. 1940. Die angioarchitektonische areale gliederung der grosshirnrinde: auf grund vollkommener gefässinjektionspräparate vom gehirn des macacus rhesus. G. Thieme.

Polioudakis D, de la Torre-Ubieta L, Langerman J, Elkins AG, Shi X, Stein JL, Vuong CK, Nichterwitz S, Gevorgian M, Opland CK, Lu D, Connell W, Ruzzo EK, Lowe JK, Hadzic T, Hinz FI, Sabri S, Lowry WE, Gerstein MB, Plath K, Geschwind DH. 2019. A Single-Cell Transcriptomic Atlas of Human Neocortical Development during Mid-gestation. Neuron 103:785–801.e8.

Rakic P. 1988. Specification of cerebral cortical areas. Science 241:170–176.

Rakic P, Ayoub AE, Breunig JJ, Dominguez MH. 2009. Decision by division: making cortical maps. Trends Neurosci 32:291–301.

Resnik P. 1995. Using Information Content to Evaluate Semantic Similarity in a Taxonomy. arXiv [cmp-lg].

Robinson EC, Garcia K, Glasser MF, Chen Z, Coalson TS, Makropoulos A, Bozek J, Wright R, Schuh A, Webster M, Hutter J, Price A, Cordero Grande L, Hughes E, Tusor N, Bayly PV, Van Essen DC, Smith SM, Edwards AD, Hajnal J, Jenkinson M, Glocker B, Rueckert D. 2018. Multimodal surface matching with higher-order smoothness constraints. Neuroimage 167:453–465.

Romero-Garcia R, Whitaker KJ, Váša F, Seidlitz J, Shinn M, Fonagy P, Dolan RJ, Jones PB, Goodyer IM, NSPN Consortium, Bullmore ET, Vértes PE. 2018. Structural covariance networks are coupled to expression of genes enriched in supragranular layers of the human cortex. Neuroimage 171:256–267.

Ronan L, Fletcher PC. 2015. From genes to folds: a review of cortical gyrification theory. Brain Struct Funct 220:2475–2483.

Ronan L, Voets N, Rua C, Alexander-Bloch A, Hough M, Mackay C, Crow TJ, James A, Giedd JN, Fletcher PC. 2014. Differential tangential expansion as a mechanism for cortical gyrification. Cereb Cortex 24:2219–2228.

Rubin TN, Koyejo O, Gorgolewski KJ, Jones MN, Poldrack RA, Yarkoni T. 2017. Decoding brain activity using a large-scale probabilistic functional-anatomical atlas of human cognition. PLoS Comput Biol 13:e1005649.

Ruzicka B, Mohammadi S, Davila-Velderrain J, Subburaju S, Tso R, Hourihan M, Kellis M. 2021. Single-Cell Dissection of Schizophrenia Reveals Neurodevelopmental-Synaptic Link and Transcriptional Resilience Associated Cellular State. Biol Psychiatry 89:S106.

Ruzzo EK, Pérez-Cano L, Jung J-Y, Wang L-K, Kashef-Haghighi D, Hartl C, Singh C, Xu J, Hoekstra JN, Leventhal O, Leppä VM, Gandal MJ, Paskov K, Stockham N, Polioudakis D, Lowe JK, Prober DA, Geschwind DH, Wall DP. 2019. Inherited and De Novo Genetic Risk for Autism Impacts Shared Networks. Cell 178:850–866.e26.

Satterstrom FK, Kosmicki JA, Wang J, Breen MS, De Rubeis S, An J-Y, Peng M, Collins R, Grove J, Klei L, Stevens C, Reichert J, Mulhern MS, Artomov M, Gerges S, Sheppard B, Xu X, Bhaduri A, Norman U, Brand H, Schwartz G, Nguyen R, Guerrero EE, Dias C, Autism Sequencing Consortium, iPSYCH-Broad Consortium, Betancur C, Cook EH, Gallagher L, Gill M, Sutcliffe JS, Thurm A, Zwick ME, Børglum AD, State MW, Cicek AE, Talkowski ME, Cutler DJ, Devlin B, Sanders SJ, Roeder K, Daly MJ, Buxbaum JD. 2020. Large-Scale Exome Sequencing Study Implicates Both Developmental and Functional Changes in the Neurobiology of Autism. Cell 180:568–584.e23.

Schaefer A, Kong R, Gordon EM, Laumann TO, Zuo X-N, Holmes AJ, Eickhoff SB, Yeo BTT. 2018. Local-Global Parcellation of the Human Cerebral Cortex from Intrinsic Functional Connectivity MRI. Cereb Cortex 28:3095–3114.

Seidlitz J, Nadig A, Liu S, Bethlehem RAI, Vértes PE, Morgan SE, Váša F, Romero-Garcia R, Lalonde FM, Clasen LS, Blumenthal JD, Paquola C, Bernhardt B, Wagstyl K, Polioudakis D, de la Torre-Ubieta L, Geschwind DH, Han JC, Lee NR, Murphy DG, Bullmore ET, Raznahan A. 2020. Transcriptomic and cellular decoding of regional brain vulnerability to neurogenetic disorders. Nat Commun 11:3358.

Singh T, Poterba T, Curtis D, Akil H, Al Eissa M, Barchas JD, Bass N, Bigdeli TB, Breen G, Bromet EJ, Buckley PF, Bunney WE, Bybjerg-Grauholm J, Byerley WF, Chapman SB, Chen WJ, Churchhouse C, Craddock N, Curtis C, Cusick CM, DeLisi L, Dodge S, Escamilla MA, Eskelinen S, Fanous AH, Faraone SV, Fiorentino A, Francioli L, Gabriel SB, Gage D, Gagliano Taliun SA, Ganna A, Genovese G, Glahn DC, Grove J, Hall M-H, Hamalainen E, Heyne HO, Holi M, Hougaard DM, Howrigan DP, Huang H, Hwu H-G, Kahn RS, Kang HM, Karczewski K, Kirov G, Knowles JA, Lee FS, Lehrer DS, Lescai F, Malaspina D, Marder SR, McCarroll SA, Medeiros H, Milani L, Morley CP, Morris DW, Mortensen PB, Myers RM, Nordentoft M, Olivares AM, Ongur D, Ouwehand WH, Palmer DS, Paunio T, Quested D, Rapaport MH, Rees E, Rollins B, Kyle Satterstrom F, Schatzberg A, Scolnick E, Scott L, Sharp SI, Sklar P, Smoller JW, Sobell J l., Solomonson M, Stevens CR, Suvisaari J, Tiao G, Watson SJ, Watts NA, Blackwood DH, Borglum A, Cohen BM, Corvin AP, Esko T, Freimer NB, Glatt SJ, Hultman CM, McQuillin A, Palotie A, Pato CN, Pato MT, Pulver AE, St. Clair D, Tsuang MT, Vawter MP, Walters JT, Werge T, Ophoff RA, Sullivan PF, Owen MJ, Boehnke M, Neale BM, Daly MJ. 2020. Exome sequencing identifies rare coding variants in 10 genes which confer substantial risk for schizophrenia. medRxiv 2020.09.18.20192815.

Sjöstedt E, Zhong W, Fagerberg L, Karlsson M, Mitsios N, Adori C, Oksvold P, Edfors F, Limiszewska A, Hikmet F, Huang J, Du Y, Lin L, Dong Z, Yang L, Liu X, Jiang H, Xu X, Wang J, Yang H, Bolund L, Mardinoglu A, Zhang C, von Feilitzen K, Lindskog C, Pontén F, Luo Y, Hökfelt T, Uhlén M, Mulder J. 2020. An atlas of the protein-coding genes in the human, pig, and mouse brain. Science 367. doi:10.1126/science.aay5947

Smith SM, Douaud G, Chen W, Hanayik T, Alfaro-Almagro F, Sharp K, Elliott LT. 2021. An expanded set of genome-wide association studies of brain imaging phenotypes in UK Biobank. Nat Neurosci 24:737–745.

Spocter MA, Hopkins WD, Barks SK, Bianchi S, Hehmeyer AE, Anderson SM, Stimpson CD, Fobbs AJ, Hof PR, Sherwood CC. 2012. Neuropil distribution in the cerebral cortex differs between humans and chimpanzees. J Comp Neurol 520:2917–2929.

Szklarczyk D, Gable AL, Lyon D, Junge A, Wyder S, Huerta-Cepas J, Simonovic M, Doncheva NT, Morris JH, Bork P, Jensen LJ, Mering C von. 2019. STRING v11: protein-protein association networks with increased coverage, supporting functional discovery in genome-wide experimental datasets. Nucleic Acids Res 47:D607–D613.

Tam V, Patel N, Turcotte M, Bossé Y, Paré G, Meyre D. 2019. Benefits and limitations of genome-wide association studies. Nature Reviews Genetics. doi:10.1038/s41576-019-0127-1

Tasic B, Menon V, Nguyen TN, Kim TK, Jarsky T, Yao Z, Levi B, Gray LT, Sorensen SA, Dolbeare T, Bertagnolli D, Goldy J, Shapovalova N, Parry S, Lee C, Smith K, Bernard A, Madisen L, Sunkin SM, Hawrylycz M, Koch C, Zeng H. 2016. Adult mouse cortical cell taxonomy revealed by single cell transcriptomics. Nat Neurosci 19:335–346.

Toro R, Burnod Y. 2005. A Morphogenetic Model for the Development of Cortical Convolutions. Cereb Cortex 15:1900–1913.

Van Essen DC. 2020. A 2020 view of tension-based cortical morphogenesis. Proc Natl Acad Sci U S A. doi:10.1073/pnas.2016830117

Velmeshev D, Schirmer L, Jung D, Haeussler M, Perez Y, Mayer S, Bhaduri A, Goyal N, Rowitch DH, Kriegstein AR. 2019. Single-cell genomics identifies cell type-specific molecular changes in autism. Science 364:685–689.

von Economo CF, Koskinas GN. 1925. Die cytoarchitektonik der hirnrinde des erwachsenen menschen. J. Springer.

Vos de Wael R, Benkarim O, Paquola C, Lariviere S, Royer J, Tavakol S, Xu T, Hong S-J, Langs G, Valk S, Misic B, Milham M, Margulies D, Smallwood J, Bernhardt BC. 2020. BrainSpace: a toolbox for the analysis of macroscale gradients in neuroimaging and connectomics datasets. Commun Biol 3:103.

Wagstyl K, Larocque S, Cucurull G, Lepage C, Cohen JP, Bludau S, Palomero-Gallagher N, Lewis LB, Funck T, Spitzer H, Dickscheid T, Fletcher PC, Romero A, Zilles K, Amunts K, Bengio Y, Evans AC. 2020. BigBrain 3D atlas of cortical layers: Cortical and laminar thickness gradients diverge in sensory and motor cortices. PLoS Biol 18:e3000678.

Weinstein SM, Vandekar SN, Adebimpe A, Tapera TM, Robert-Fitzgerald T, Gur RC, Gur RE, Raznahan A, Satterthwaite TD, Alexander-Bloch AF, Shinohara RT. 2021. A simple permutation-based test of intermodal correspondence. Hum Brain Mapp 42:5175–5187.

Werling DM, Pochareddy S, Choi J, An J-Y, Sheppard B, Peng M, Li Z, Dastmalchi C, Santpere G, Sousa AMM, Tebbenkamp ATN, Kaur N, Gulden FO, Breen MS, Liang L, Gilson MC, Zhao X, Dong S, Klei L, Cicek AE, Buxbaum JD, Adle-Biassette H, Thomas J-L, Aldinger KA, O’Day DR, Glass IA, Zaitlen NA, Talkowski ME, Roeder K, State MW, Devlin B, Sanders SJ, Sestan N. 2020. Whole-Genome and RNA Sequencing Reveal Variation and Transcriptomic Coordination in the Developing Human Prefrontal Cortex. Cell Rep 31:107489.

Xia J, Zhang C, Wang F, Meng Y, Wu Z, Wang L, Lin W, Shen D, Li G. 2018. A COMPUTATIONAL METHOD FOR LONGITUDINAL MAPPING OF ORIENTATION-SPECIFIC EXPANSION OF CORTICAL SURFACE AREA IN INFANTS. Proc IEEE Int Symp Biomed Imaging 2018:683–686.

Xu X, Sun C, Sun J, Shi W, Shen Y, Zhao R, Luo W, Li M, Wang G, Wu D. 2022. Spatiotemporal Atlas of the Fetal Brain Depicts Cortical Developmental Gradient. J Neurosci 42:9435–9449.

Yarkoni T, Poldrack RA, Nichols TE, Van Essen DC, Wager TD. 2011. Large-scale automated synthesis of human functional neuroimaging data. Nat Methods 8:665–670.

Yeo BTT, Krienen FM, Sepulcre J, Sabuncu MR, Lashkari D, Hollinshead M, Roffman JL, Smoller JW, Zöllei L, Polimeni JR, Fischl B, Liu H, Buckner RL. 2011. The organization of the human cerebral cortex estimated by intrinsic functional connectivity. J Neurophysiol 106:1125–1165.

Yip AM, Horvath S. 2007. Gene network interconnectedness and the generalized topological overlap measure. BMC Bioinformatics 8:22.

Zeng H, Shen EH, Hohmann JG, Oh SW, Bernard A, Royall JJ, Glattfelder KJ, Sunkin SM, Morris JA, Guillozet-Bongaarts AL, Smith KA, Ebbert AJ, Swanson B, Kuan L, Page DT, Overly CC, Lein ES, Hawrylycz MJ, Hof PR, Hyde TM, Kleinman JE, Jones AR. 2012. Large-scale cellular-resolution gene profiling in human neocortex reveals species-specific molecular signatures. Cell 149:483–496.

Zhang B, Horvath S. 2005. A general framework for weighted gene co-expression network analysis. Stat Appl Genet Mol Biol 4:Article17.

Zhang Y, Sloan SA, Clarke LE, Caneda C, Plaza CA, Blumenthal PD, Vogel H, Steinberg GK, Edwards MSB, Li G, Duncan JA 3rd, Cheshier SH, Shuer LM, Chang EF, Grant GA, Gephart MGH, Barres BA. 2016. Purification and Characterization of Progenitor and Mature Human Astrocytes Reveals Transcriptional and Functional Differences with Mouse. Neuron 89:37–53.

