## Supplementary figures for "Transcriptional Cartography Integrates Multiscale Biology of the Human Cortex"

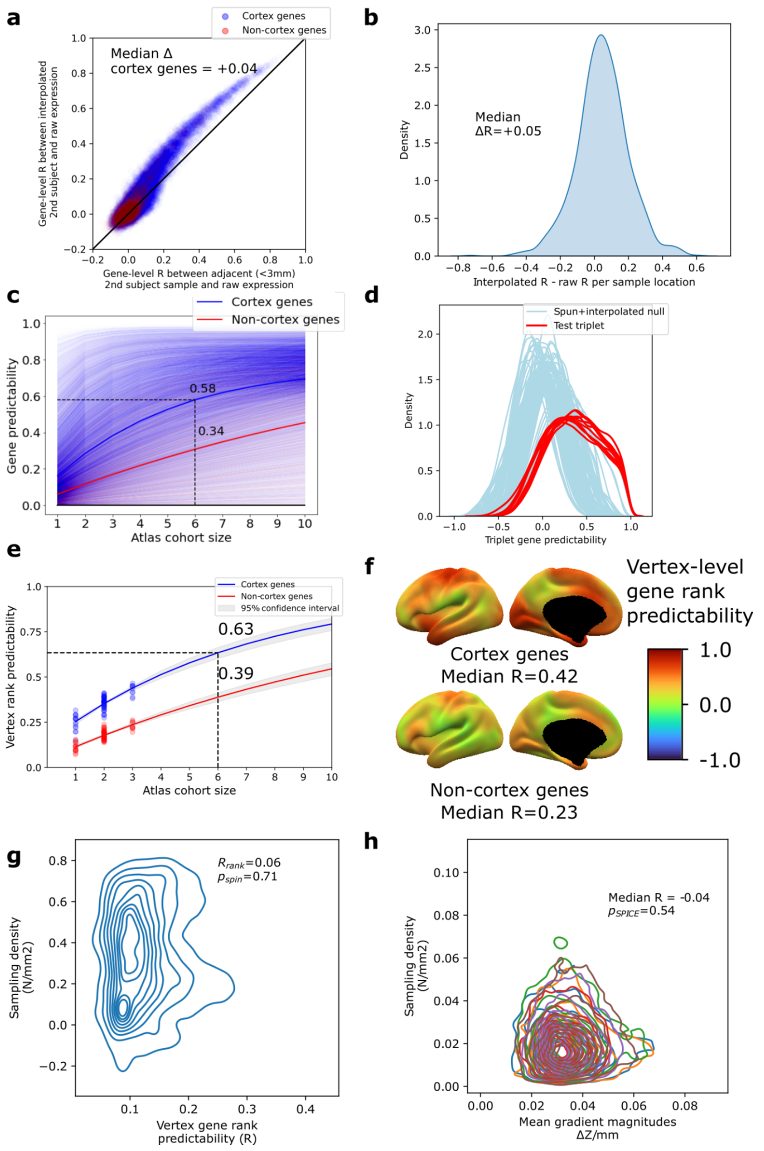


***Figure S1. Reproducibility of Dense Expression Maps (DEMs) interpolated from spatially sparse postmortem measures of cortical gene expression.*** ***a,*** *Signal boost in the interpolated DEM dataset vs. spatially sparse expression data. Restricting to samples taken from approximately the same cortical location in pairs of individuals (within 3mm geodesic distance), there was an overall improvement in intersubject spatial predictability in the interpolated maps. Furthermore, genes with lower predictability in the interpolated maps were less predictable in the raw dataset, suggesting these genes exhibit higher underlying biological variability rather than methodologically introduced bias.* ***b,*** *Similarly at the paired sample locations, gene-rank predictability was generally improved in DEMs vs. sparse expression data (median change in R from sparse samples to interpolated data for each pair of subjects, +0.5).* ***c,*** *Gene-level reproducibility of spatial patterns from DEMs independently generated by subsampling the 6 donor cohort into groups of 1, 2 and 3 subjects (median across splits). Learning curve analyses of median gene DEM predictability using DEMs created with subsamples of the full cohort, were generated separately for cortically expressed and non-cortical genes. The spatial reproducibility of DEMs increases with the number of sample donors and can be extrapolated to estimate reproducibility using the full sample size of 6, giving an estimated r of 0.58.* ***d,*** *Gene predictability was higher across all triplet-triplet pairs than when compared to the spun+interpolated null.* ***e,*** *Vertex-level reproducibility of gene rankings by scaled expression at each cortical vertex calculated for using the subsampling processes as above. Learning curves were fit to median per vertex predictability of relative gene ranks with different sample cohort sizes. Using the full sample size of 6, cortically*

expressed genes had an estimated reproducibility of 0.63. ***f,*** *Ranking of genes by scaled expression at each cortical vertex is generally reproducible and more so for cortically expressed vs. non-cortically expressed genes.* ***g,*** *The magnitudes of gradients of DEMs were not associated with the local sampling density.* ***h,*** *Comparison of the sampling density with the reliability of gene expression rankings showed no relationship.*

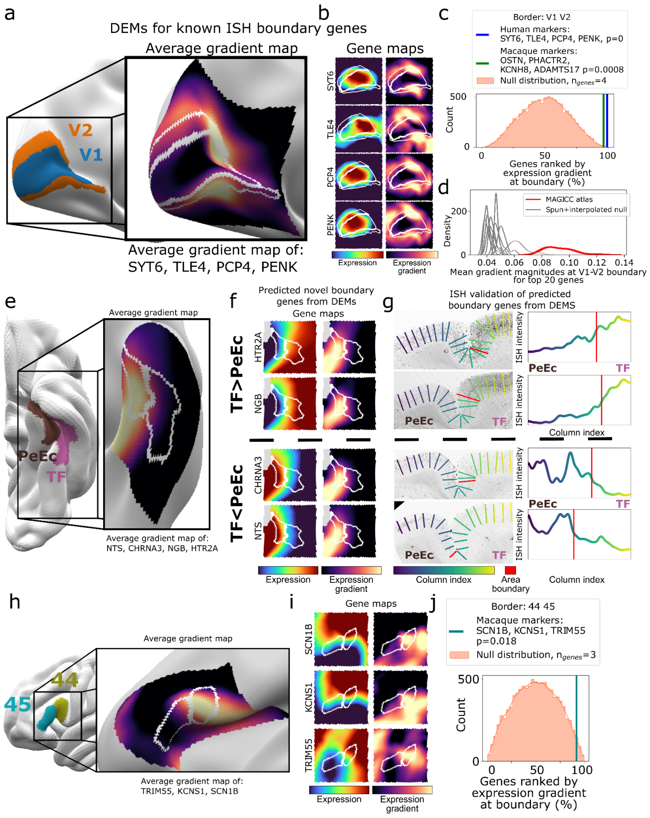


*Figure S2* ***Validating and discovering area marker genes.*** ***a,*** *Validation local DEM gradients through areal marker genes. Gene expression gradients were averaged along the border between V1 and V2, shown on an inflated cortical surface. The average of four recognised V1 area markers (Zeng et al., 2012 Cell) exhibits high changes in expression along the boundary between V1 and V2.* ***b,*** *Each marker gene exhibits a clear peak of expression within V1 with high expression gradients in mm  outside the region.* ***c****, Mean gradient for known marker genes at the V1 V2 border in both macaques angmd humans were significantly highly ranked relative to randomly sampled groups of genes.* ***d,*** *Mean of gradient magnitudes for 20 genes with largest gradients along V1-V2 border, compared to values along the same boundary on the spun+interpolated null atlas. Gradients were higher in the actual dataset than in all spun version indicating this high gradient feature is not primarily due to the effects of calcarine sulcus morphology on interpolation.* ***e,*** *DEM gradients used for discovery of novel area markers. Quantified along the sulcal border between areas PeEc (parahippocampal gyrus) and TF (fusiform gyrus),* ***f,*** *Each of the four DEM gradient genes exhibited clear expression changes along this border.* ***g,*** *Putative border genes were validated using available ISH data capturing the same dividing sulcus. These genes exhibited visible and quantifiable changes in expression in the predicted location.* ***h,*** *DEM gradients in relatively cytoarchitectonically homogeneous frontal regions, 44 and 45, mark areal boundary genes identified in the macaque.* ***i,*** *DEM gradients for known marker genes at the area 44 / 45 border in macaque were significantly highly ranked relative to randomly sampled groups of genes.*

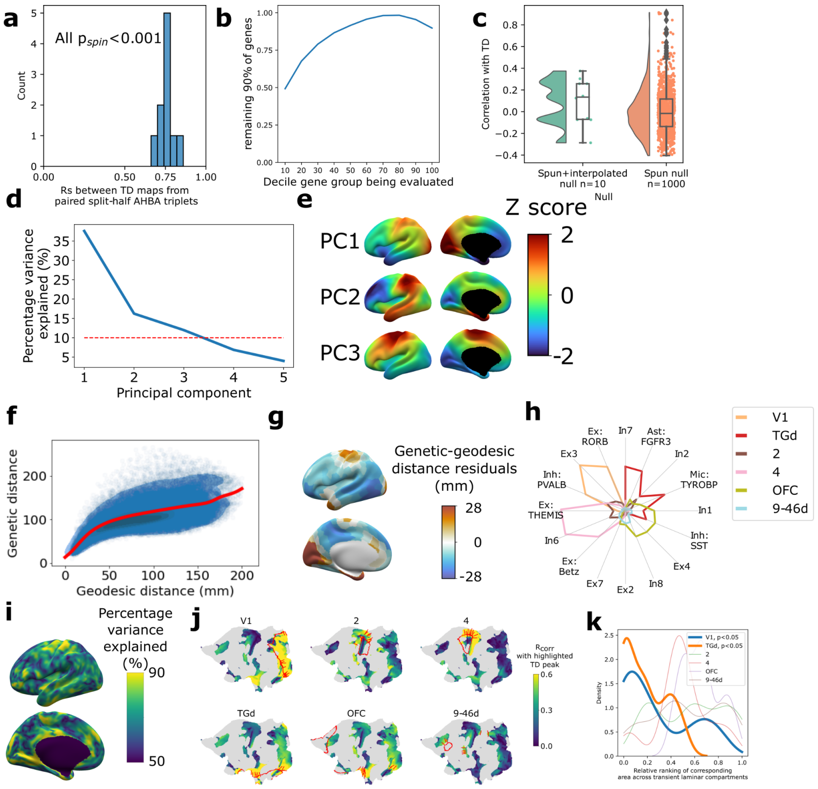


***Figure S3. Characterizing bulk transcriptome. a,*** *Triplet reproducibility of cortical Transcriptomic Distinctiveness (TD). Across all ten combinations of triplets from the 6-subject cohort, the median correlation between pairs of triplets was r=0.77,* $p_{spin}$*<0.001.* ***b,*** *Genes were grouped into deciles according to the reproducibility of their spatial patterns in independent sub-cohorts (Fig S1c). Each decile's TD map was compared to the map from the remaining 9 deciles.* ***c,*** *Correlations between TD and TD maps regenerated on datasets spun using two independent nulls, one where the rotation is applied prior to interpolation and smoothing (spun+interpolated) and one where it is applied to the already-created DEMs. In each null, the same rotation matrix is applied to all genes.* ***d,*** *Percentage variance explained by the first five principal components of cortical gene expression, compared to a spatially permuted null (red-dashed line).* ***e,*** *DEMs of statistically significant first 3 PCs, shown on an inflated cortical surface.* ***f,*** *Pairwise inter-regional transcriptomic distances against geodesic cortical distances. Red line shows the prediction from a Generalized Additive Model.* ***g,*** *Median residual transcriptomic distances for each region, characterizing areas that are transcriptomically further from other areas than their cortical physical embedding would predict.* ***h,*** *Cortical cell type overlaps with TD peak regions. 21 of 24 DEMs for cortical cell marker genes had peaks located in TD peak regions (*$p_{spin}$*<0.001), with TD peaks showing distinctive overlaps.* ***i,*** *Percentage variance explained by the first principal component of the gene expression gradient vectors. High percentages indicate concerted, anisotropic gene expression gradients.* ***j,*** *Flattened representations of the cortical sheet with areas of significantly high magnitude gradients. Vertex colors are given by rank correlation of gene gradient magnitudes with gene ranks at the labeled TD peak (outlined). The principal orientation of gene expression gradients are shown in red for selected high magnitude vertices, and can be seen to run perpendicular to TD peak boundaries.* ***k,*** *The mean expression z-score of high-ranking genes in each TD peak was calculated for measures of gene expression in the fetal cortex at 21 PCW, sampled at six transient fetal layers with an average of 29 regional samples per layer. Because these regional measures of fetal expression sampled multiple cortical regions at different transient fetal layers, we could rank regions within a layer by their expression of genes characterizing a given TD peak and then ask if high-ranking fetal regions for a given TD tended to include regions overlapping that TD (i.e. do fetal visual cortex samples tend to be amongst the highest ranked for V1 TD marker genes from adult DEMs?). These relative intra-layer rankings are shown as density plots for each TD, with bold lines for those two adult TDs (V1 and TGd) that already have localized transcriptional identities at 21 PCW.*

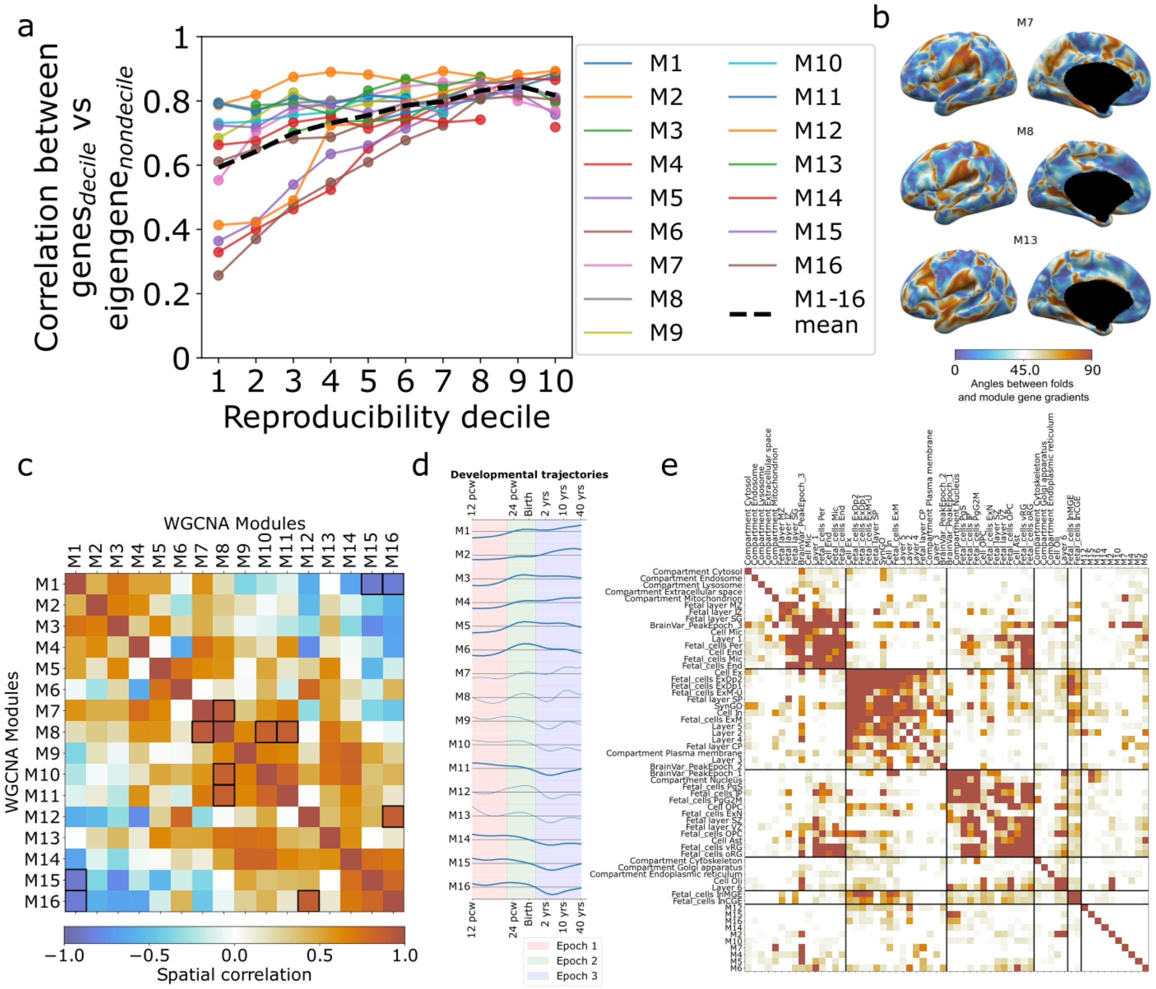

***Figure S4. Characterization of WGCNA spatial, developmental and gene-set relationships*** ***a,*** *Gene DEMs for each module were stratified into deciles according to their reproducibility score and correlated with their module's eigengene. Higher reproducibility genes tended to resemble the module eigengene more closely.* ***b,*** *Strength of cortical folding alignment with WGCNA modules M7, M8 and M13.* ***c,*** *Spatial correlations between WGCNA module eigenmaps. Significant correlations after controlling for multiple comparisons are outlined in black.* ***d,*** *Average developmental trajectory of genes in modules. In bold are modules where gene-level trajectories exhibit significantly greater intra-modular correlation than expected by chance.* ***e,*** *Matrix representation of significant pairwise overlaps between annotational gene sets used in* ***Fig 3c****, including WGCNA module gene sets and a UMAP embedding of the matrix is presented in* ***Fig 3e****. Black lines demarcate clusters of annotations.*

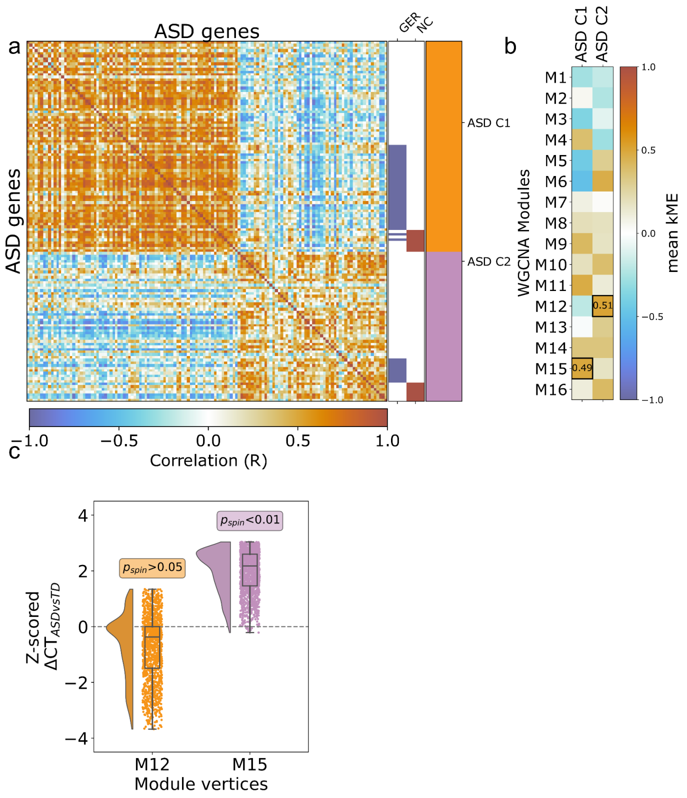


***Figure S5. a,*** *Clustering of DEMs associated with ASD risk genes identified two contrasting clusters, ASD C1 & ASD C2. C1 was primarily enriched for genes associated with the Gene Expression Regulation, with C2 for Neuronal Communication.* ***b,*** *Consistent with findings from* ***Fig 4****, across all 16 WGCNA modules, DEMs for genes in C1 had the highest mean kME with M15 eigenmap, while genes in C2 had the highest mean kME with M12.* ***c,*** *The region of peak M15 expression (but not M12 expression) shows significantly greater cortical thickening in ASD than other cortical regions.*
