## Supplemental tables for "Transcriptional Cartography Integrates Multiscale Biology of the Human Cortex"

**Supplementary Table 1. Study participants demographics**

1. Demographics of adult donors Allen Human Brain Atlas microarray data
2. Demographics of fetal samples from Allen Institute's fetal Laser Microarray Dataset
3. Demographics of included participants for Alzheimer’s disease APOE analysis from the Open Access Series of Imaging Studies (OASIS)
4. Demographics of included participants for Autism Spectrum Disorder analysis from the Autism Brain Imaging Data Exchange (ABIDE) I & II.

**Supplementary Table 2. Gene lists used in the study**

1. Gene list assignments for enrichment analyses including WGCNA modules.
2. Meta module assignments

**Supplementary Table 3. Transcriptomically distinctive (TD) peaks**

1. TD genes, GO, cellular, fetal and functional annotations
2. Remaining sheets describe significant Biological Process and Cellular Compartment Gene Ontology annotations for TD peaks

**Supplementary Table 4. WGCNA module enrichments**

1. WGCNA module spatial and gene set enrichment p-values.
2. Remaining sheets describe significant Biological Process and Cellular Compartment Gene Ontology annotations for WGCNA modules
